# PARP1 promotes replication-independent DNA double-strand break formation after acute DNA-methylation damage

**DOI:** 10.1101/2025.07.10.663928

**Authors:** Anne Marie McMahon, Haichao Zhao, Jia Li, Garrett Driscoll, Joshua Matos, Kelly McGhee, Jade Lyttle, Shan Yan

## Abstract

Poly-ADP-Ribose Polymerase 1 (PARP1) is a potent regulator of DNA damage response signaling through the recruitment of DNA damage repair proteins to damage sites, and its catalytic function of converting Nicotinamide adenine dinucleotide (NAD^+^) into poly-ADP-ribose (PAR) which covalently modifies hundreds of protein substrates in a process known as PARylation. However, PARP1’s role in the recognition, processing, and intracellular signaling downstream of DNA damage in cells remains incompletely understood, especially in a replication-independent context. Here, we show that cells exposed to high doses of the methylating agent Methyl Methanesulfonate (MMS) generate DNA double-strand breaks (DSBs) in a base excision repair (BER)-dependent and DNA replication-independent manner. The capacity of cells to generate DSBs after MMS exposure relies heavily on intracellular NAD^+^ availability and PARP1’s catalytic production of PAR. In our experimental system, we show that acute MMS exposure causes NAD^+^ exhaustion in a PARP1-dependent manner, which results in a temporal-dependent loss of downstream PARP1 activity. This functional loss of PARP1 signaling in later timepoints leads to the loss of BER-dependent single-strand break (SSB)-to-DSB conversion, as well as silencing of ATR-Chk1 signaling in both cycling and non-cycling cells, demonstrating a novel PARP1-dependent regulatory mechanism for both ATR-Chk1 signaling and BER-associated processes following methylation challenge. Additionally, we provide experimental evidence supporting the role of PARP1 and NAD^+^ in promoting the exonuclease-mediated SSB-to-DSB conversion. These findings support a previously uncharacterized mechanism of PARP1-mediated replication-independent DSB generation and provide insight into checkpoint signaling by integrating DDR with PARP1’s consumption of NAD^+^ and production of PAR.

## INTRODUCTION

Methyl Methanesulfonate (MMS) is a mono-functional alkylating agent that primarily methylates DNA at N^7^-guanine (N^7^-meG, ∼80%), with minor modifications at N^3^-adenine (N^3^-meA, ∼10%) and O^6^-guanine (O^6^-meG, <0.5%)^1,2^. Importantly, the predominant lesion generated by MMS (N^7^- meG) is not thought to be DNA polymerase stalling and is therefore “hidden” within the helix and not sensed by the DNA damage response (DDR) until it is converted into DNA repair intermediates such as abasic (AP) sites, single strand breaks (SSB), or double strand breaks (DSB) by DNA repair pathways such as base excision repair (BER)^3–5^. Among these intermediates, DSBs are considered the most genotoxic due to their capacity to trigger apoptosis and cause chromosomal rearrangements^6^. For over five decades, the mechanism by which MMS exposure results in DSBs in the genome has been debated, with two predominant models proposed. First, the DNA replication stress model suggests that DSBs arise from stalled DNA replication forks, caused by DNA polymerase collision with N^3^-meA, O^6^-meG, or BER intermediates, leading to single-ended DSBs^7–9^. Secondly, the SSB-to-DSB conversion model proposes that alkylated bases are converted into SSBs and that two proximal SSBs on opposite strands will be converted into one DSB^10,11^. It is proposed that these proximal SSBs must be within one helical turn to cause this spontaneous DSB conversion^12^, although this remains to be experimentally validated under different conditions (GC content, helical tension, etc.). Notably, the SSB-to-DSB conversion model is the same proposed mechanism through which ionizing radiation (IR) is thought to generate DSBs and may explain why MMS- and IR-induced DNA damage in cells rely on many of the same DNA repair and signaling pathways. This similarity to IR has led to MMS being historically characterized as a ‘radiomimetic’ agent^13–20^. While MMS-induced DNA replication fork stalling and DSB formation following MMS exposure has been appreciated for nearly 50 years by experimental evidence, the proximal SSB-to-DSB conversion model, to the best of our knowledge, has not been directly supported by experimental data^4,12,21–25^. This, coupled with the distance constraints on SSB-to-DSB conversion, has cast serious doubt on its plausibility^12^. In this study, we present direct evidence of DNA replication-independent DSBs induced by MMS, supporting the existence of SSB-to-DSB conversion after MMS treatment. Furthermore, we propose that a potent protein sensor of DNA damage, Poly-ADP-Ribose Polymerase 1 (PARP1), plays a key role in directing DNA replication-independent DSB generation, as well as DDR signaling, after acute MMS challenge.

PARP1, a major regulator of the DNA damage response and DNA damage repair environments, plays an important but obscure role in the recognition and repair of DSBs and SSBs^26^. PARP1 possesses the ability to reprogram protein-protein interactions through the generation of multimeric chains of poly-ADP-ribose (PAR), which can modify local environments such as biomolecular condensates, through direct and indirect mechanisms of action^27–30^. Because of these properties, PARP1 lies at several critical junctions between different cellular processes such as DSB repair pathway choice^31–33^, metabolism and cell fate^34–36^. PARP1 converts its substrate NAD^+^ into PAR, a polymer similar RNA^26^. PARP1’s use of NAD^+^ as a substrate plays a significant role in connecting metabolism and DNA repair; however, the role of the PARP1/NAD^+^ junction has not been shown to regulate the DDR signaling significantly.

Three phosphatidylinositol-3-kinase–related protein kinases named Ataxia-telangiectasia mutated (ATM), Ataxia-telangiectasia and RAD3-related (ATR), and DNA-dependent protein kinase catalytic subunit (DNA-PKcs) regulate DSB repair and repair pathway choice^37^. These DDR kinases are known to phosphorylate hundreds of substrates, including themselves and each other, which is determinative of multiple cellular fates, including cell cycle arrest via Chk1 and Chk2 proteins, apoptosis and transcriptional reprograming through p53^38–40^. Of these three “master regulator” kinases, ATR is arguably the most sensitive, as it is known to be activated by recruitment to RPA-bound single-strand DNA (ssDNA)^41,42^. RPA-ssDNA can be generated in all cell cycle phases. While ATR signaling is best known for its roles in DNA replication stress, it has been observed to be activated by ssDNA generation in a replication-independent manner, likely through exonuclease processing of SSBs^43–46^.

Here, we demonstrate that NAD^+^ availability and PARP1-mediated PAR generation are key determinants of two key cellular outcomes after acute MMS challenge: DSB generation and ATR-Chk1 signaling. Within ∼1 hour of treatment, PARP1 rapidly depletes NAD^+^, and by 2–3 hours, cells lose the capacity to generate nascent PAR. This functional loss of PARP1 activity correlates with a loss of exonuclease activity, which we propose is necessary for converting BER intermediates into DSBs by reducing the spacing constraints between nearby SSBs. Moreover, the diminished exonuclease activity may underlie the observed reduction in ATR–Chk1 signaling, reducing the amount of RPA-coated ssDNA substrate required for ATR activation. Together, our findings suggest a novel role for PARP1 in regulating both ATR signaling maintenance and exonuclease processing of BER intermediates. They also underscore the importance of NAD^+^ availability in maintaining DDR capacity and cellular resilience following genotoxic stress, particularly from agents like IR, which generates DNA lesions through similar clustered damage mechanisms^47^.

## RESULTS

### MMS exposure generates DSBs in a BER-dependent manner in cells

MMS is a widely used genotoxic agent in DNA damage repair studies and is known to generate DSBs in yeast and human cells. Here, we confirm that acute MMS exposure (3mM) is capable of generating DSBs after a one-hour exposure time (Fig. 1A-1E). To accurately assess DSB counts per cell, we labeled the DSB marker protein and upstream promoter of NHEJ, Tumor suppressor p53-binding protein 1 (53BP1), as well as the phosphorylated histone H2AX at Serine 139, γH2AX, a DNA damage marker not exclusive to DSBs^48–52^. Both of these proteins form distinct puncta/foci which are easily identified and counted via immunofluorescent microscopy^53^. Importantly, only when these foci colocalize/overlap are they counted as a DSB for the purposes of this study. Here we reproducibly observed an average of 15 colocalized 53BP1-γH2AX foci per nuclei after one hour of MMS exposure in U2OS cells (Fig. 1A-1B). Surprisingly, 53BP1 foci are virtually undetectable by two- and three-hour MMS treatment (Fig. 1A-1B). γH2AX foci numbers were observed to increase incrementally from no treatment until one-hour MMS exposure, while no significant change was found from one hour to three hours (Fig. 1A and 1C). To alleviate concerns about the half-life of MMS, we prepared fresh 3mM MMS containing media and replaced the MMS media every hour to ensure that cells undergo active alkylation damage for the whole duration of the 3h window (Fig. S1D-S1F). This media swap control did not affect the 53BP1 formation and dissipation dynamics compared with the non-swapped media experiments (Fig. S1D-S1F), demonstrating that potential MMS depletion was not contributing to our observations. Importantly, while we perform most experiments here in U2OS cells, we are also able to reproduce the DSB formation and resolution dynamics in PANC1 cells with and without MMS media swap conditions (Fig. S1G-S1J).

**Figure 1.**
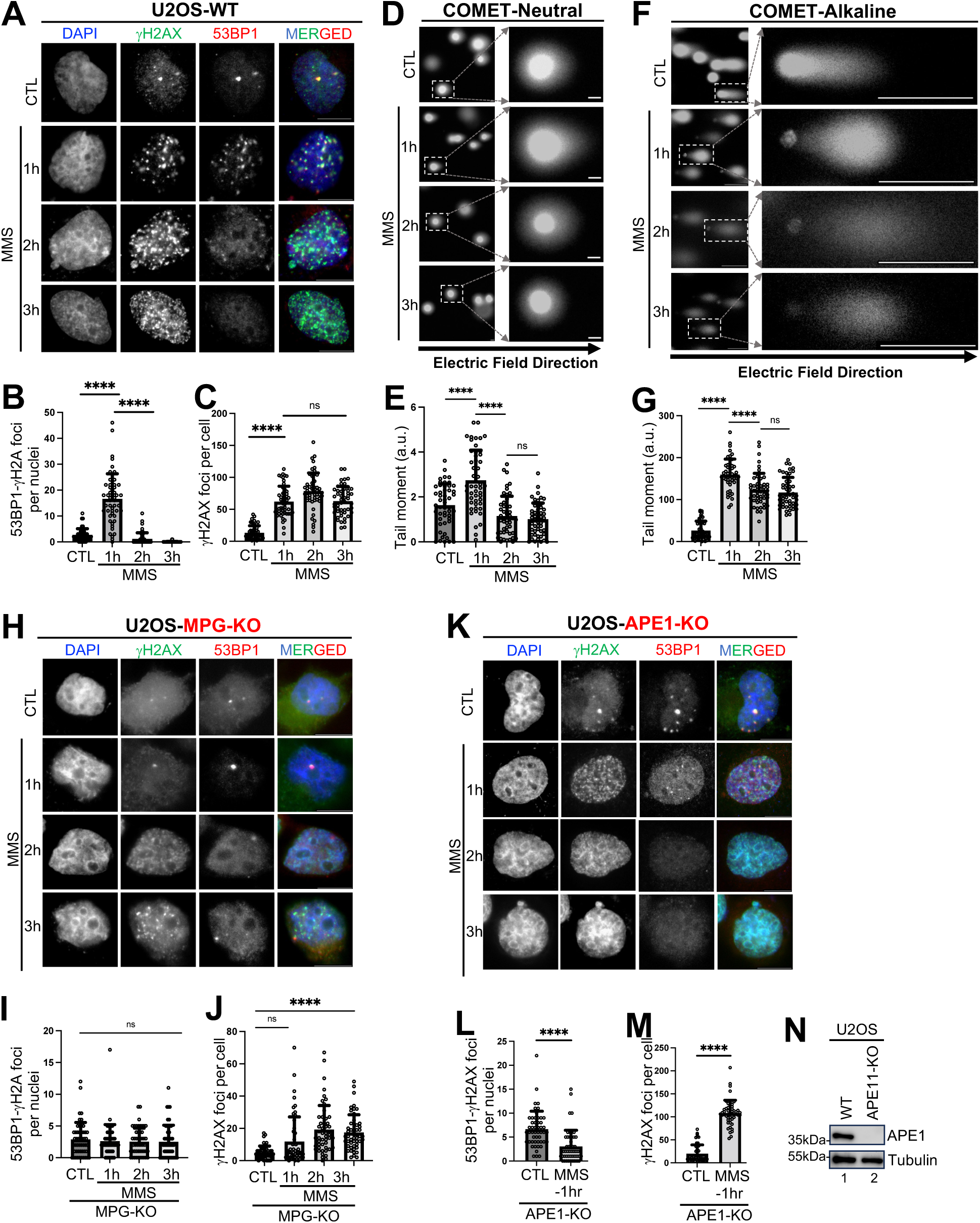
MMS generates DSB in a BER-dependent manner. *A*, U2OS cells were used for immunofluorescent microscopy using DAPI (nuclei staining), γH2AX (AF488), and 53BP1 (AF594) after exposing cells to one, two, and three hours of 3mM MMS respectively. *B*, Overlapping/colocalizing γH2AX-53BP1 foci were quantified on a per nuclei basis using n=50 cells. *C*, γH2AX foci per cell were quantified for n=50 cells after MMS exposure. *D*, Comet assay under neutral condition was performed after indicated 3mM MMS exposure. *E,* Tail moment was quantified using Comet Assay IV Lite software for n=50 cells from *D*. *F*, Comet assay under alkaline condition was performed after indicated 3mM MMS exposure. *G*, Tail moment was quantified using Comet Assay IV Lite software for n-=50 cells from *F*. *H*, U2OS MPG-KO cells were used for immunofluorescent microscopy using DAPI (nuclei staining), γH2AX (AF488), and 53BP1 (AF594) after exposing cells to one, two, and three hours of 3mM MMS respectively. *I*, Overlapping/colocalizing γH2AX-53BP1 foci were quantified on a per nuclei basis using n=50 cells from *H*. *J*, γH2AX foci per nuclei were quantified for n=50 cells from *H*. *K*, U2OS APE1-KO cells were used for immunofluorescent microscopy using DAPI (nuclei staining), γH2AX (AF488), and 53BP1 (AF594) after exposing cells to one, two, and three hours of 3mM MMS respectively. *L* Overlapping/colocalizing γH2AX-53BP1 foci were quantified on a per nuclei basis using n=50 cells from *K*. *M*, γH2AX foci per nuclei were quantified for n=50 cells from *K*. *N*, Immunoblot confirmation of APE1-KO cells was performed in parallel with WT U2OS cells. For all experiments, the Kolmogorov-Smirnov test was used to determine statistical significance and is demonstrated as follows: ****, p<0.0001; ns, no significance. Scale bar = 10μm.

Two possible explanations for 53BP1 foci resolution are (1) the DSBs are repaired and the mechanism driving DSB formation is turned off, or (2) 53BP1 is removed from DSB ends through DSB end processing and/or remodeling. To interrogate these possible models, we performed the comet assay under neutral (DSB detecting) and alkaline (all lesion detecting) conditions^54^ and found that neutral comet produced tails which aligned with 53BP1 recruitment and dissociation dynamics (Fig. 1D-1E). This data strongly suggests that DSBs are generated rapidly within 1h of MMS treatment and are rapidly resolved under continuous MMS challenge. Total DNA damage loads measured via alkaline comet show a slight reduction in tail moment between one and two hours, which corresponds with our window of DSB repair (Fig. 1F-1G).

Additionally, we performed quantitative intensity analysis of γH2AX-stained cells and found that the intensity of γH2AX increased dramatically over time, where γH2AX intensity is roughly 100x higher in three-hour-treated samples compared with one-hour-treated samples (Fig. S1A-S1C). This, coupled with consistently high tails in alkaline comet (Fig. 1F-1G), demonstrates that MMS causes increased “global” DDR activation throughout the duration of the treatment, likely via the generation of non-DSB DNA damage such as SSBs, ssDNA gaps, transcription stress, replication stress, and others.

Nitrogen moieties of purine rings are the most common target of environmental alkylating agents and also of MMS^55^. Removal of *N*-alkylpurines is catalyzed by BER machinery which starts with the DNA glycosylase *N*-methylpurine-DNA glycosylase (AAG/MPG) to generate AP sites. AP sites are converted into SSBs by AP Endonuclease 1 (APE1). SSBs are sensed by PARP1, which is thought to promote additional downstream BER protein recruitment, such as Ligase 3 (Lig3) and DNA Polymerase β (Polβ), for repair synthesis and ligation^56^. MMS lesions must be converted from *N*-alkylpurines into proximal SSBs on opposite strands via the BER pathway to support a proximal SSB-to-DSB conversion mechanism. Exonuclease processing of SSBs towards an adjacent SSB may also increase the likelihood of a DSB conversion taking place. If SSB-to-DSB conversion is a major mechanism of MMS-induced DSB, MPG and APE1 would be obligate upstream effectors of MMS-generated DSB.

Consistent with the SSB-to-DSB conversion model, we find that MPG-knockout (KO) and APE1-KO U2OS cell lines did not produce DSB after MMS exposure (Fig. 1H-1N). Compared with WT cells, MPG-KO cells had reduced γH2AX foci (Fig. 1H, 1J), supporting the idea that MMS lesions do not activate DDR until BER processing. APE1-KO resulted in surprisingly high numbers of γH2AX foci (Fig. 1M), likely because of unrepaired AP site collisions with DNA and RNA polymerases. However, neither MPG-KO nor APE1-KO cells produced significant 53BP1-γH2AX foci following MMS exposure (Fig. 1H-1N). In addition, we performed siRNA-mediated knockdown of MPG and APE1 and, similar to our knockout conditions, observed reduced 53BP1-γH2AX foci at 1-hour MMS treatment in MPG-knockdown (KD) and APE1-KD U2OS cells, compared with control knockdown U2OS cells (Fig. S2A-S2C, S2E-S2F). These observations suggest that MPG and APE1 are both important for MMS-induced DSB generation, consistent with the idea that downstream BER processing of MMS-induced base damage is required for DSB generation.

Taken together, this data suggests that dynamic intracellular conditions dictate the DSB generation capacity of MMS, possibly through regulation of BER activity. Our findings here agree with previous experiments which have shown that MMS-induced methyl-base repair intermediates generated by MPG are a driving force in DSB generation^9^. We expand this idea by investigating the role of APE1 in the generation of DSB after MMS and find APE1 is also critical for the DSB formation capacity of MMS challenge, highlighting the importance of AP-site to SSB conversion for MMS-induced DSB.

### NAD^+^ availability and PARP1-dependent PAR production promote MMS-induced DSB formation

NAD^+^ is an essential metabolite that is best known for its roles in central metabolism and nucleotide biosynthesis. NAD^+^ is also consumed as a substrate for PARP1 in the catalytic generation of PAR. PARP1 thus bridges metabolism and the DDR by modulating the available pools of NAD^+^ in response to DNA damage. PARP1’s consumption of NAD^+^ is also thought to play a major role in PARP1-mediated cell death signaling. To measure PARP1 activity and NAD^+^ availability, antibody-mediated detection of PAR can be used for both immunoblotting and immunofluorescent analysis. Here, we show that PAR generation and dissociation dynamics reflect the patten of DSB formation after MMS exposure, where one-hour MMS treatment induces robust PAR generation followed by a decline in PAR signal at two- and three-hour marks (Fig. 2A). PARP1-KO cells do not mount a significant PAR response to MMS challenge, demonstrating that MMS-induced PAR generation is mainly PARP1-dependent (Fig. 2A).

**Figure 2.**
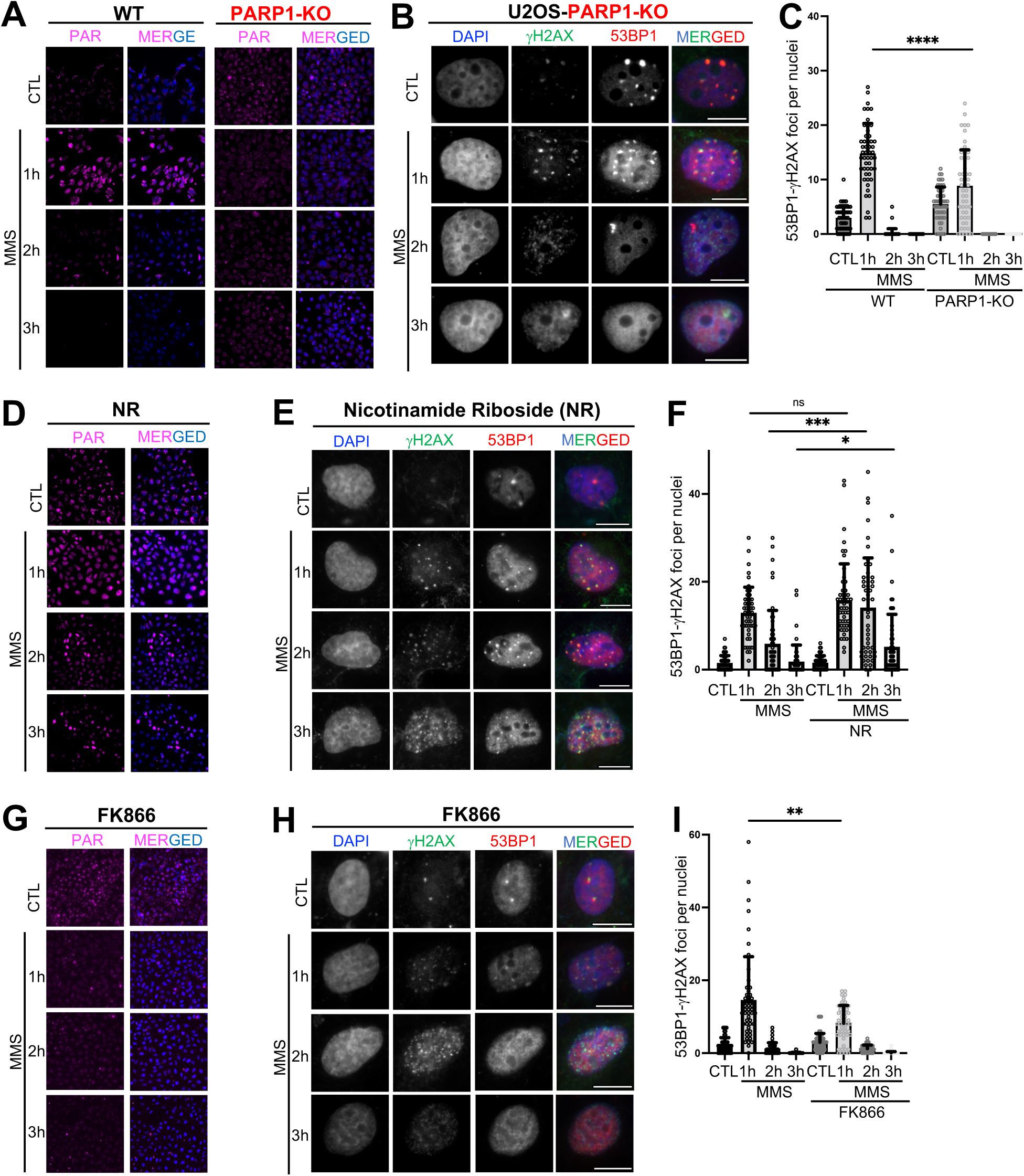
NAD^+^ availability and PARP1-dependent PAR production promote MMS-induced DSB formation. *A*, PAR (magenta) was imaged using immunofluorescence (AF488) and DAPI (nuclei) staining after indicated exposure to 3mM MMS in both WT and PARP1-KO U2OS cells. *B*, U2OS PARP1-KO cells were treated with MMS in parallel with WT U2OS cells and stained for DAPI, 53BP1 (AF594) and γH2AX (AF488). *C*, Overlapping/colocalizing γH2AX-53BP1 foci were quantified on a per nuclei basis using n=50 cells from *B*. *D*, PAR was imaged after supplementation with Nicotinamide Riboside Chloride (NR) (200μM for 24 hours pre-treatment) and subsequent MMS challenge. *E*, NR treatment was performed before immunofluorescent microscopy using DAPI (nuclei staining), γH2AX (AF488), and 53BP1 (AF594) after exposing cells to one, two, and three hours of 3mM MMS respectively in parallel with untreated controls. *F*, Overlapping/colocalizing γH2AX-53BP1 foci were quantified on a per nuclei basis using n=50 cells from *E*. *G*, PAR was imaged after treatment with FK866 (50μM for 24 hours pre-treatment) and MMS challenge as indicated. *H* FK866 treatment was performed before immunofluorescent microscopy using DAPI (nuclei staining), γH2AX (AF488), and 53BP1 (AF594) after exposing cells to one, two, and three hours of 3mM MMS, respectively, in parallel with untreated controls. *I* Overlapping/colocalizing γH2AX-53BP1 foci were quantified on a per nuclei basis using n=50 cells from *H*. For all experiments, Kolmogorov-Smirnov test was used to determine statistical significance, and statistical significance is demonstrated as follows: *, p<0.05; **, p<0.01; ***, p<0.001; ****, p<0.0001; ns, no significance. Scale bar = 10μm.

We also tested the effect of PARP1-KO on the capacity for DSB generation. PARP1-KO cells showed a significant reduction in initial DSB formation (∼9 DSB in PARP1-KO vs ∼15 in WT), supporting the idea that PARP1 may contribute to DSB by generating PAR in response to MMS treatment (Fig. 2B-2C, and S3A). Consistent with the PARP1-KO observation, we performed siRNA-mediated PARP1-KD before staining for 53BP1-γH2AX foci and observed reduced DSB generation and γH2AX intensity in PARP1-KD cells, compared with control knockdown cells (Fig. S2A, S2E-S2G).

To better explore the idea that NAD^+^ availability, and thus cellular capacity to generate PAR, may modulate MMS-induced DSB formation, we established an NAD^+^ supplement regimen by adding Nicotinamide Riboside Chloride (NR) which could extend PAR generation into the two- and three-hour timepoints (Fig. 2D). To reduce intracellular NAD^+^, we use the NAMPT inhibitor FK866 which was effective at reducing PAR response to MMS challenge after one hour (Fig. 2G), providing us with both a strategy to enhance PARP1 activity and nascent PAR production into the later MMS timepoints (NR treatment) and a strategy to diminish PARP1 activity at early timepoints compared with untreated controls (FK866 treatment). In addition to this, we use PARG inhibitor (PARGi) PDD00017273^57,58^, which prevents degradation of PAR produced in response to MMS, but notably it does not extend the ability of PARP1 to produce nascent PAR after NAD^+^ pools are depleted (Fig. S3E-S3F).

Using NR and FK866 to increase and decrease intracellular NAD^+^ levels, respectively, we tested the capacity of cells to generate DSB after MMS challenge. Notably, NAD^+^ supplementation (NR treatment) resulted in an enhanced capacity for cells to generate DSB at 2h and 3h MMS compared with untreated controls (Fig. 2E-2F). On the other hand, we found that when cells had reduced intracellular NAD^+^ levels (following FK866 treatment), they produced approximately 50% less DSB relative to untreated controls at a one-hour MMS challenge (Fig. 2H-2I). These findings strongly suggest that NAD^+^ levels directly influence cellular capacity to generate DSBs after MMS challenge. Additionally, PARGi resulted in greatly increased DSB generation at one hour MMS but did not significantly affect DSB dynamics at two- and three-hour MMS compared with untreated controls (Fig. S3C, S3E-S3F). Taken together, these results demonstrate a critical role for PAR generation and accumulation after MMS challenge in the processing of MMS-induced DNA methyl-bases.

In addition to its roles in DSB generation, PAR also appears to be critical for effective “global” DDR signaling through the γH2AX intensity response, where lowering intracellular NAD^+^ levels with FK866 treatment leads to a diminished γH2AX response (Fig. S3G-S3H), while preserving PAR via PARGi treatment results in more rapid and robust γH2AX response to MMS (Fig. S3I-S3J).

### NAD^+^ availability and PAR retention maintains ATR-Chk1 signaling

DNA damage is known to induce potent intracellular responses that facilitate the recognition, recruitment, and repair of DNA damage lesions. Interestingly, the repair of DNA damage can itself lead to genotoxic intermediates. Here, in agreement with previous studies^9^, we have demonstrated that BER is required for the generation of DSBs following MMS challenge. Several groups^9,59–61^, including us here, have shown that MMS can generate a DNA damage response as measured by γH2AX foci formation and intensity analysis. Unsurprisingly, γH2AX intensity increases in a time-dependent manner after MMS exposure (Fig. S1A-S1C). However, we have shown that MMS results in unique PAR production dynamics which appear to regulate DSB formation after MMS challenge (Fig. 2). Because of this, we wanted to explore further the downstream effects of dynamic PAR production on the DDR signaling pathways.

Our initial investigation of DDR signaling dynamics looked at ATR and DNA-PKcs kinases using well-defined DDR activation markers (ATM-P-S1981; Chk1-P-S345 and Chk1-P-S317; DNA-PKcs-P-S2056). We observed that MMS challenge induces robust ATR-Chk1 activation initially; however, this ATR-Chk1 signaling returns to baseline levels after just three hours MMS exposure (Fig. 3A-3B). Once ATR-Chk1 signaling begins to decline, DNA-PKcs signaling turns on (Fig. 3A-3B). We initially believed that this signaling “handoff” may indicate a direct role of ATR in the preclusion of DNA-PKcs signaling after DSB formation. However, we later identified several conditions (Fig. 3E-3J) which allowed the manipulation of ATR-Chk1 signaling during the three-hour MMS treatment but did not significantly alter DNA-PKcs activation dynamics. Because of this, we tentatively propose that the activation dynamics of DNA-PKcs are more a result of initial DSB generation at one hour after MMS and subsequent repair than due to a preclusive role of ATR signaling. This idea is also supported by the concept that DNA-PKcs-P-S2056 may indicate completed NHEJ repair of DSBs rather than initial DSB recognition^62–67^.

**Figure 3.**
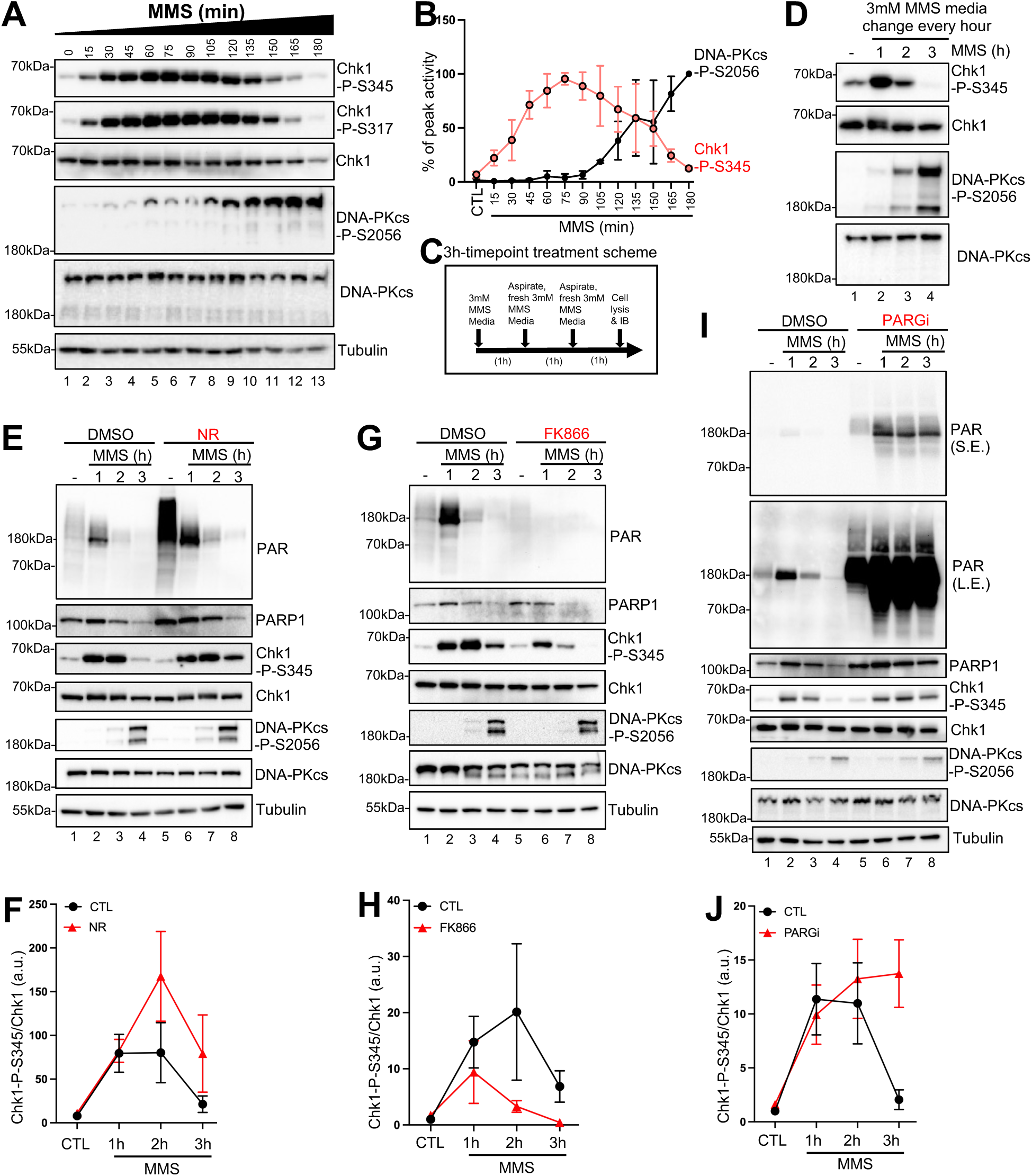
NAD^+^ availability and PAR retention maintains ATR-Chk1 signaling. *A*, Immunoblotting was performed on total cell lysates after 15-minute increment samples were collected after 3mM MMS exposure, for a total of three hour of continuous MMS exposure. *B*, Percent change of Chk1-P-S345 and DNA-PKcs-P-S2056 was quantified from three independent biological replicates using ImageJ after normalization to respective total Chk1 OR DNA-PKcs signals. *C*, Three-hour treatment schema for media change experiment. *D*, Immunoblotting was performed after 3mM MMS exposure with media change every hour. *E*, NR pre-treatment was performed (200μM for 24 hours) before 3mM MMS exposure for the indicated times, followed by subsequent immunoblotting. *F*, Quantification of Chk1-P-S345 normalized to total Chk1 comparing untreated and NR-treated samples. *G*, FK866 pre-treatment was performed (50μM for 24 hours) before 3mM MMS exposure for the indicated times and subsequent immunoblotting. *H*, Quantification of Chk1-P-S345 normalized to total Chk1 comparing untreated and FK866-treated samples. *I*, PARG inhibitor pre-treatment was performed before 3mM MMS exposure for the indicated times and subsequent immunoblotting. *J*, Quantification of Chk1-P-S345 normalized to total Chk1 comparing untreated and PARGi-treated samples.

Similar to a control we performed earlier in this study, we swapped MMS-containing media every hour to confirm that the loss of ATR signaling was not due to MMS exhaustion but instead a result of dynamic intracellular changes during active MMS challenge (Fig. 3C-3D). Notably, the loss of ATR-Chk1 signaling and PAR after three hours of 3mM MMS exposure was a reproducible phenotype in PANC1 cells (Fig. S4G-S4H).

We next sought to determine if the availability of NAD^+^ and the cellular capacity to produce PAR contributed to our observed DDR signaling dynamics. We had already seen some evidence for the effect of NAD^+^ availability and PAR on “global” DDR signaling through γH2AX intensity profiling, where NAD^+^ availability and PARGi promote γH2AX formation (Fig. S3G-S3J). Here we demonstrate that reduction of intracellular NAD^+^ with FK866 treatment results in attenuated ATR-Chk1 signaling (Fig. 3G-3H) and supplementing cells with additional NAD^+^ via NR allows for extended ATR-Chk1 signaling into the three-hour MMS timepoint (Fig. 3E-3F). Importantly, when we retain PAR via PARGi, we also rescue ATR-Chk1 signaling at three hours MMS (Fig. 3I-3J). PARGi allows cells to retain most, or all intracellular PAR produced after MMS challenge but does not allow for nascent PAR production in the two- and three-hour timepoints (Fig. 3I). This suggests that it is the loss of PAR signal via PARG, and not exclusively nascent PAR production/NAD+ pool depletion that mechanistically results in the loss of ATR-Chk1 signaling after three-hour MMS treatment. The exact mechanism of ATR-Chk1 signaling retention via PAR remains unknown; however, we uncovered a critical role of PARP1 in promoting exonuclease activity on SSBs later in this study (Fig. 4). Exonuclease activity on DNA breaks generates stretches of ssDNA for RPA association to promote the ATR-Chk1 DDR signaling activation^41,43,44,68^. We suggest that cellular capacity for exonuclease processing of SSB (and also DSB) may contribute to our observed phenotypes. This idea is explored further in subsequent sections.

**Figure 4.**
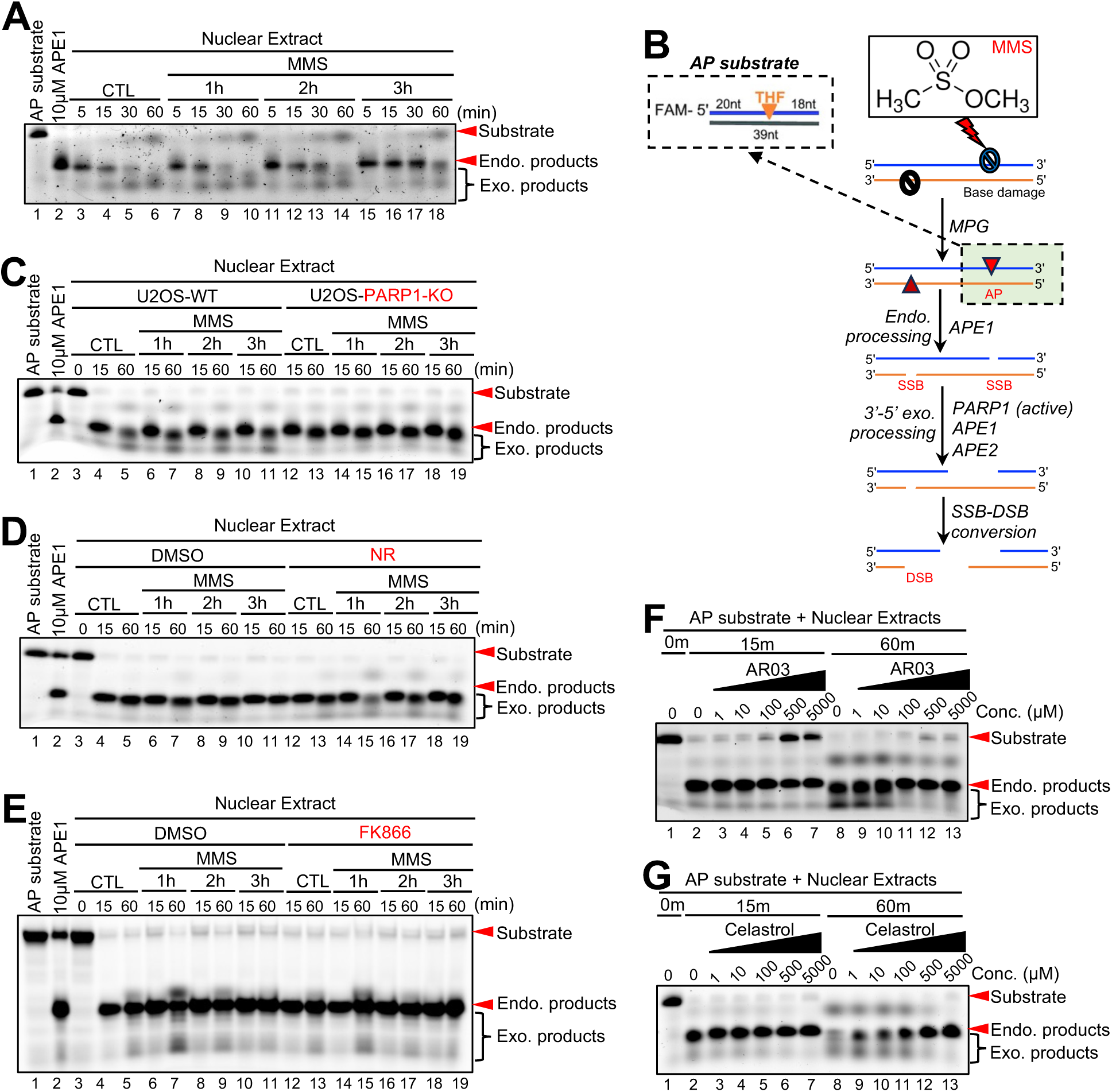
PARP1, APE1, and APE2 promote the exonuclease activity of nuclear extracts following MMS challenge. *A*, Nuclear extract was prepared from cells after MMS exposure for indicated times. 20μg of nuclear extract was incubated with the DNA construct for indicated times at 37°C, followed by denaturing EMSA assay. *B*, FAM-labeled AP-site mimic construct and SSB-to-DSB model. *C*, Nuclear extracts were prepared from WT and PARP1-KO U2OS cells after indicated MMS exposures. 20 μg of nuclear extract was incubated with the DNA construct for the indicated times at 37°C, followed by denaturing EMSA assay. *D*, Nuclear extracts were prepared from untreated and NR-treated WT U2OS cells (200μM for 24 hours pre-treatment) after indicated MMS exposures. 20 μg of nuclear extract was incubated with the DNA construct for the indicated times at 37°C, followed by a denaturing EMSA assay. *E*, Nuclear extracts were prepared from untreated and FK866-treated WT U2OS cells (50μM for 24 hours pre-treatment) after indicated MMS exposures. 20 μg of nuclear extract was incubated with the DNA construct at 37°C for the indicated times, followed by denaturing EMSA assay. *F*, Nuclear extracts were prepared from WT U2OS cells. 20 μg of nuclear extract was incubated with indicated concentrations of AR03 for 30 minutes prior to the addition of the DNA construct for indicated times at 37°C, followed by denaturing EMSA assay. *G*, Nuclear extracts were prepared from WT U2OS cells. 20 μg of nuclear extract was incubated with indicated concentrations of Celastrol for 30 minutes prior to the addition of the DNA construct for indicated times at 37°C, followed by denaturing EMSA assay.

Of note, PARP1 stability appears to be influenced by the PAR production and stability. After MMS treatment, PARP1 protein abundance is gradually lost and almost indetectable by three-hour MMS. Reduction of intracellular NAD^+^ by FK866 results in faster destabilization of PARP1 (Fig. 3G and S4A), while stabilization of PAR by PARGi results in a stabilization of PARP1 protein abundance over all timepoints (Fig. 3I and S4C). NR appears to have little to no observable effect on PARP1 protein stability (Fig. 3E and S4B). In our hands, proteasome inhibition and Caspase 3/7 inhibition did not rescue PARP1 degradation, and we did not see an observable band indicative of a PARP1 cleavage product as is seen in similar studies (data not shown)^69^. While we do not yet understand the mechanistic underpinnings of our observed PARP1 protein degradation, it is interesting that PARP1 becomes functionally inactive after MMS treatment both through exhaustion of its substrate and through loss of protein stability.

We also wanted to test the effect of MMS exposure times on cell viability under various conditions to determine which pathways may be protective or detrimental to MMS challenged cells. One hour exposure to MMS at 3mM is terminal to cells, even if recovered for more than twelve hours. To circumvent this issue and test cell viability, we designed our assay to measure cell viability after a total of six hours of recovery. Using this method, we demonstrated that, as expected, longer MMS exposure times result in a faster loss of cell viability. After looking at several pathways of interest, we found that the inhibition of PARP1 with Olaparib resulted in higher cell viability (Fig. S4E). When we supplemented the cells with NR, we achieved a similar rescue of cell viability, suggesting that PARP1 may drive cell death through consumption of intracellular energy currency (Fig. S4D). We also observed that the inhibition of DNA-PKcs via NU7441 reduced cell viability, suggesting a protective role for NHEJ in our system (Fig. S4F).

### Exonuclease activity of nuclear extracts is promoted by PARP1, APE1, and APE2

MMS treatment remodels intracellular environments and distinctly affects cell’s ability to generate DSB and activate ATR over time. PAR synthesis and the role of PARP1 in promoting BER appears to regulate these processes, however directly testing this idea is technically challenging. To better understand the mechanism underlying our observations, we developed a system in which cells are treated with MMS prior to collection and nuclear extraction. After this nuclear extraction, we incubate the nuclear extract with a defined DNA construct that mimics an AP site. We then used this AP-site mimic to compare the difference between untreated and MMS treated nuclear extracts in their efficiency of processing these DNA constructs (Fig. 4B). We found that the ability of our nuclear extracts to cleave the AP sites by APE1’s endonuclease activity does not significantly differ before and after MMS treatment (Fig. 4A). We also used different dilutions of nuclear extract as a control to demonstrate that more concentrated extract more efficiently processes the AP-site mimic (Fig. S5A).

Interestingly, we noticed that our extracts could execute 3’-5’ exonuclease activity after a longer one-hour incubation at 37°C (Fig. 4A). Initial endo-nucleolytic processing of AP-sites occurs in as little as five minutes, but a longer incubation of up to one-hour results in continued excision of the SSB into exonuclease products. Interestingly, there was a profound decrease in exonuclease activity of the nuclear extract after longer MMS exposures (Fig. 4A). Within a cellular context, this may mean exonuclease activity promotes ATR-Chk1 activation by generating longer ssDNA, which is an upstream substrate and activator of ATR. Additionally, exonuclease processing of SSB promotes SSB-to-DSB conversion by reducing the distance between proximal SSBs to within the distance needed for spontaneous DSB conversion. If exonuclease activity is diminished throughout MMS exposure, this may explain why we observe both a loss of DSB formation and a silencing of the ATR-Chk1 activation.

To test if there was any relationship between nuclear extract exonuclease activity and PARP1, we repeated these experiments with PARP1-KO cells, NR-supplemented cells, and FK866-treated cells before nuclear extraction. We find that PARP1 is essential for exonuclease processing of our AP-site construct after it has been cleaved into a SSB (Fig. 4C). Agreeing with this observation, NR-mediated supplement of NAD^+^ resulted in hyper-SSB-resection and FK866-dependent reduction of NAD^+^ pools attenuated SSB-resection via exonuclease activity (Fig. 4D-4E). These results connect our previous observations that PARP1-dependent PAR synthesis promotes the maintenance of ATR-Chk1 signaling and DSB formation after MMS challenge, offering a mechanistic underpinning for these phenotypes.

Next, we sought to identify reasonable exonuclease candidates that possess 3’-5’ exonuclease activity. Using our extracts, we tested the effects of Mre11, APE1, and APE2 inhibitors on endo- and exonuclease processing of our AP-site substrate. While we did not observe any notable exonuclease inhibition via Mre11 inhibition (data not shown), APE1 inhibitor AR03^70^ and APE2 inhibitor Celastrol^68^ both resulted in convincing dose-dependent inhibition of exonuclease processing (Fig. 4F-4G). Significantly, higher concentrations of AR03 also inhibited endonuclease cleavage of our AP-site construct, acting as an important control in our system (Fig. 4F). While we previously showed that knockout and knockdown of APE1 resulted in a decrease in DSB formation after MMS challenge (Fig. 1K-1N, S2C, S2E), APE2’s role in DSB formation was unknown. We tested the effect of siRNA-mediated APE2-KD or APE2 inhibition by Celastrol on DSB formation. While siRNA-mediated APE2-KD had no appreciable impact on DSB formation at 1 hour MMS exposure, APE2 inhibition via Celastrol resulted in apparent reduction of DSB formation after MMS challenge (Fig. S5B-S5F). The differential effect of APE2-KD and APE2 inhibition in MMS-induced DSB formation may be due to several factors, such as a compensatory exonuclease, which may become functional in the absence of APE2 protein. Because previous studies have shown the role of APE2 in HR and MMEJ repair of DSBs^71,72^, we can’t exclude the scenario that the reduced DSB repair capacity of cells in the absence of APE2 may complicate the potential reduction of DSB generation by APE2 absence. We also must consider that while our knockdown appears to be highly effective via western blot (Fig. S5B), a small amount of APE2 protein may be present that is below our detection limit, which could be confusing our results. It should also be noted that Celastrol is known to have other protein targets (e.g., HSP90) and off-target effects cannot be excluded as a contribution to our observed phenotype^73,74^. We cautiously propose that Celastrol inhibits exonuclease activity by acting on its previously described exonuclease target, APE2^68^, resulting in the attenuation of DSB formation. What is clear is that Celastrol robustly inhibits exonuclease processing of our substrate *in vitro* and strongly reduces DSB formation *in vivo*, supporting the importance of exonuclease activity on DSB formation after MMS challenge.

### PARP1 promotes DSB formation in a DNA replication-independent manner

MMS can induce a broad spectrum of DNA methyl adducts in cells including N^7^-meG (∼80%) and N^3^-meA (∼10%) as major base damages, and O^6^-methylated guanine (O^6^-meG), N^1^-methyladenine (N^1^-meA) and N^3^-methylcytocine (N^3^-meC) as minor base lesions^1^. While the minor base damages are repaired by MGMT-mediated direct reversal of methyl group or ALKBH2/ALKBH3-mediated demethylation repair pathway (Fig. S6A)^1,75–78^, the major base damages (N^7^-meG and N^3^-meA) can be processed into AP sites by N-methylpurine DNA glycosylase MPG (also known as AAG) and subsequently converted into SSBs by APE1 (Fig. S6A)^1,9,77,79^. The DNA replication stress model suggests that one-ended DSBs are generated when DNA replication forks collide with BER intermediates, such as SSBs (Fig. S6A)^8,9,80^. The SSB-DSB conversion model proposes that proximal SSBs on opposite strands will be converted into DSBs containing 3’-ssDNA overhang, 5’-ssDNA overhang, or blunt ends (Fig. S6A)^10,11,81^. While the replication-dependent formation of DSB has been appreciated for some time, we decided to explore the possibility that MMS may be able to produce DSB in a replication-independent manner.

We postulate that if MMS only generates DSBs in S-phase then we expect to see roughly 30-50% of cells with DSB (representing cells in S-phase) and 50-70% of cells without DSB (representing G1 & G2 phase cells). However, when we observed cell populations using our immunofluorescent analysis, this bias was not evident in asynchronous cells. To visualize this, we plotted DSB counts from our one-hour MMS-treated cells and made QQ plots and frequency distributions of this data set. We compared our distributions to a simulated data set, assuming 50% of the cell population is in S-phase, an average of 20 DSBs per cell affected, and a standard deviation of 10 DSBs in affected cells (Fig. S6B). The simulated counts show the bimodal distribution that would be expected for a cell-cycle specific DSB-inducing drug (Fig. S6C-S6D). In contrast, our observed DSB distribution is more “normal” and shows a positive skew distribution (Fig. S6E-S6F). Because of this, we predicted that MMS may not exhibit a substantial bias between cell cycle phases, and we hypothesized that MMS may be capable of generating DSBs in an S-phase-independent manner via an exonuclease-mediated SSB-to-DSB conversion mechanism.

To test this idea experimentally, we synchronized cells in G1 phase of the cell cycle using a previously established Thymidine-Nocodazole synchronization protocol^82^, and then treated them with MMS to determine if DSBs were still generated. In agreement with the SSB-to-DSB conversion model, we observed the robust generation of DSB after MMS treatment both through the generation of 53BP1-γH2AX colocalized foci formation (Fig. 5A-5C) and the neutral comet assay (Fig. 5D-5E) in the synchronized G1 phase cells. The quantification of these foci only revealed a slight decline in the average number of 53BP1-γH2AX colocalized foci in comparison with asynchronous cells (after 1h MMS: ∼10/cell in G1 phase cells vs ∼15/cell in asynchronous cells), suggesting that the SSB-to-DSB conversion model significantly contributes to, but is not the only factor, in the generation of DSB after acute MMS challenge to an asynchronous population of cells (Fig. 5B). In addition to this approach, we also performed EdU labelling of replicating S-phase cells from asynchronous cell populations using click chemistry to label genome-incorporated EdU with Alexa Fluor 488. Using this approach, we can visualize 53BP1 foci formation in non-replicating cells (G1 & G2 phase cells), which represent a large majority of high-count 53BP1 foci positive cells (Fig. 5F-5H). Importantly, whether cells are replicating or not, DSB generation appears to be effectively turned off after one hour of MMS exposure, and the cell cycle state seems to not affect the capacity of cells to resolve these MMS-induced DSBs (Fig. 5A-5E). Taken together, these results provide the first clear evidence for replication-independent DSB formation after MMS treatment.

**Figure 5.**
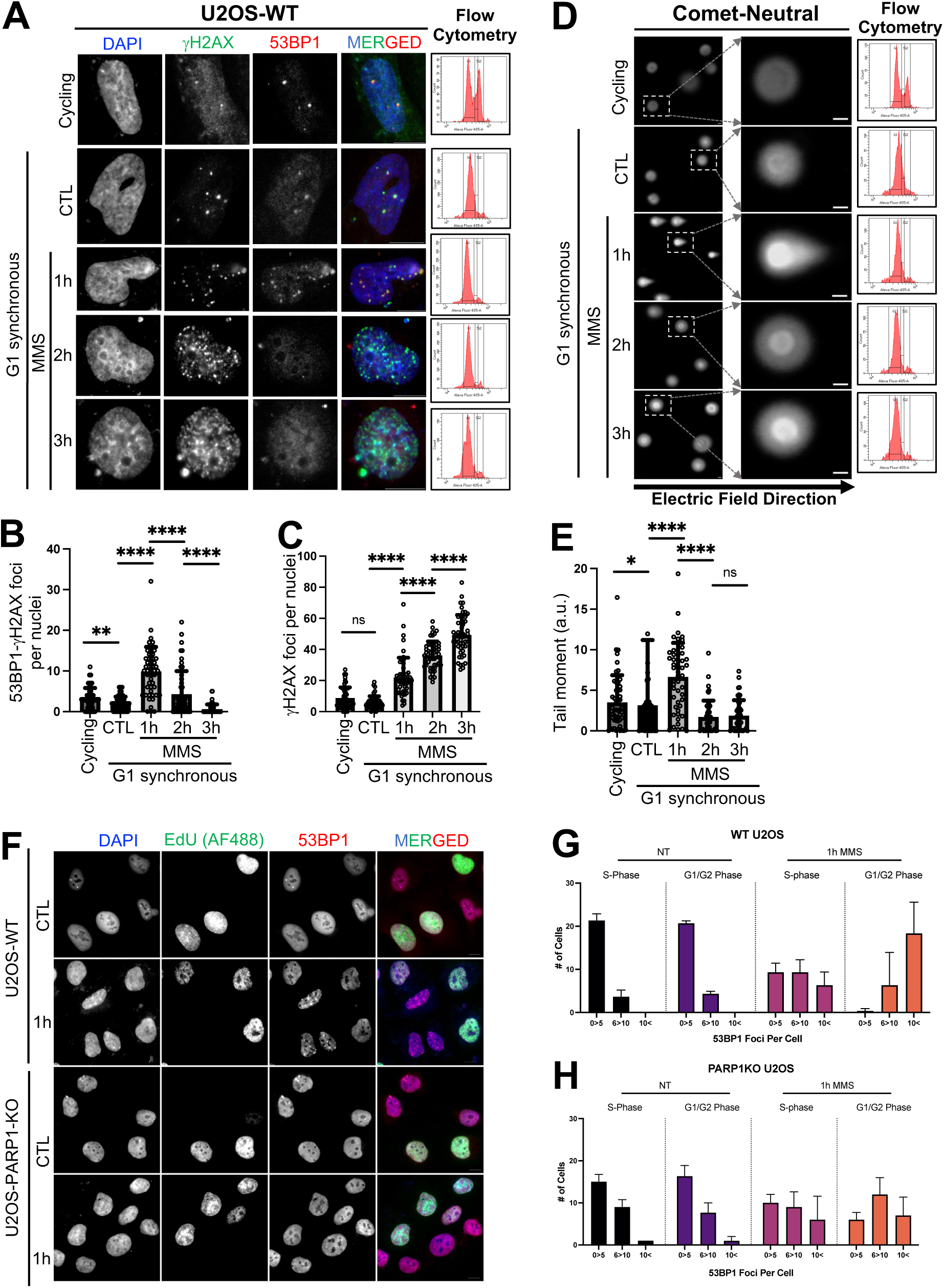
PARP1 promotes DSB formation in a DNA replication-independent manner. *A*, U2OS WT cells were synchronized in G1 phase for immunofluorescent microscopy using DAPI (nuclei staining), γH2AX (AF488), and 53BP1 (AF594) after exposing cells to one, two, and three hours of 3mM MMS respectively. *B*, Overlapping/colocalizing γH2AX-53BP1 foci were quantified on a per nuclei basis using n=50 cells from *A*. *C*, γH2AX foci per nuclei were quantified for n=50 cells from *A*. *D*, U2OS WT cells were synchronized to G1 phase before neutral Comet assay was performed after indicated 3mM MMS exposure. *E,* Tail moment was quantified using Comet Assay IV Lite software for n=50 cells from *D*. *F,* EdU labelling was performed on WT and PARP1-KO U2OS cells for 2 hours prior to one hour of MMS exposure as indicated. DAPI, EdU (AF488), and 53BP1 (AF594) were labeled and imaged under appropriate channels. *G*, WT U2OS quantification for 53BP1 foci per cell, stratified based on cell cycle stage (n=25 cells were quantified from 3 biological replicates for a total n=75 cells, bars represent the mean of these three replicates and error bars represent standard deviation between replicates). *H*, PARP1-KO U2OS quantification for 53BP1 foci per cell, stratified based on cell cycle stage (n=25 cells were quantified from 3 biological replicates for a total n=75 cells, bars represent the mean of these three replicates and error bars represent standard deviation between replicates). For all experiments, the Kolmogorov-Smirnov test was used to determine statistical significance and is demonstrated as follows: *, p<0.05; **, p<0.01; ****, p<0.0001; ns, no significance. Scale bar = 10μm.

Based on PARP1’s importance for exonuclease activity in our nuclear extract system, we predicted that PARP1 loss may cause a reduction of DSB formation primarily in non-replicating cells. To test this, we labelled wild-type and PARP1-KO asynchronous cells with EdU to identify cells in S-phase. We then stained for 53BP1 foci and stratified our counts based on cell cycle (S-phase vs G1/G2 phase). We found that WT cells and PARP1-KO cells share similar distributions of 53BP1 foci formation in S-phase cells, but PARP1-KO cells have markedly fewer 53BP1 foci in G1/G2 phase cells compared with WT U2OS (Fig. 5F-5H). This data provides cellular-level evidence for the importance of PARP1 in promoting SSB-to-DSB conversion, linking our anticipated cellular phenotype with reduced SSB exonuclease processing in PARP1-KO cells (Fig. 4).

We also synchronized our cells to G1 phase to see if there was any S-phase requirement for ATR-Chk1 activation and subsequent silencing and found no difference between G1 phase and asynchronous cells, indicating that ATR-Chk1 activation and subsequent silencing, similar to DSB formation, is DNA replication-independent (Fig. S6G). This supports the idea that loss of exonuclease-mediated generation of ssDNA in later MMS timepoints is a contributing factor to the silence of the ATR-Chk1 signaling.

In this study, we show that DSBs are generated from SSBs which are a downstream intermediate of BER processing of MMS-induced methylation damage. The accumulation and processing of these SSBs results in the activation of the ATR-mediated DNA damage response and the production of DSBs in a DNA replication-independent, and exonuclease-dependent process which is regulated by PARP1 and the production of PAR. Our study provides evidence to support the idea that the biochemical pathways responsible for the recognition and repair of DNA damage can participate in the production of deleterious repair intermediates. Overall, our findings highlight the critical molecular mechanism of PARP1, NAD^+^, and PARylation in the regulation of DSB generation following radiomimetic MMS exposure via the SSB-to-DSB conversion model in mammalian cells.

## DISCUSSION

In this study we demonstrate that DSBs are generated rapidly after MMS exposure in a BER-dependent but replication-independent manner. PARP1-mediated synthesis of PAR promotes this DSB formation, likely by promoting exonuclease-mediated processing of SSBs which allows for spontaneous SSB-to-DSB conversion.

Initially, we hypothesized that prolonged MMS exposure would lead to a time-dependent increase in number of DSBs. Contrary to our expectation, we saw that MMS generates DSB rapidly (∼1 hour) and loses its capacity to generate DSBs in cells after a longer exposure (2- and 3-hours). We confirmed this observation both through 53BP1 foci imaging and via comet assay-based experiments. Excitingly, we found that MMS can generate DSBs in a replication-independent, BER-dependent manner (Fig. 1 & 5). This claim is backed by evidence in G1-phase synchronized cells and by EdU labeling in asynchronous cell populations (Fig. 5). Previous publications emphasize DSB formation by MMS during S-phase, which certainly occurs, but ignore the potential capacity of MMS-induced replication-independent DSB formation. Indeed, we believe our findings point to a broader flaw in genotoxin-agent-based studies where a well-established damaging agent in question (such as MMS here) is known to produce a phenotype (replication fork collapse), but interpretations of the data can be confounded by additional phenotypes (replication-independent DSB). Better characterization of genotoxic agents’ mechanisms of action will continually improve our interpretation of subsequent data. Interestingly, replication-independent DNA damage goes against many of the conventional ideas of chemotherapy. Many chemotherapies target cell’s DNA replication pathways based on the well-supported notion that cancer cells have upregulated DNA replication and downregulated DNA repair. While MMS is not currently used as a chemotherapy drug, it is possible that many of these currently used chemotherapy agents also induce DNA damage and DSB in G1 & G2 phases of the cell cycle through poorly understood mechanisms. A better understanding of cell-cycle dependencies could allow for improved clinical application of genotoxic chemotherapies.

MMS-induced DSB heavily relies on NAD^+^ availability and PARP1-mediated synthesis of PAR. Extension or premature termination of PARP1 activity with NAD^+^ supplement (NR) or NAMPT inhibitor (FK866) respectively, resulted in tunable changes in MMS’s capacity to generate DSB, where supplementation of NAD^+^ resulted in more DSB generation capacity in the later timepoints of MMS exposure, and reduction of NAD^+^ pools diminished MMS-induced DSB levels. Based on these observations, we propose that PARP1 promotes BER-dependent DSB formation through its catalytic function (Fig. 3).

The silencing of ATR-Chk1 signaling (after initial activation) also appears to depend on a loss of PAR signal after one hour of MMS. This loss of ATR-Chk1 signaling represents, to the best of our knowledge, the first instance where this signaling pathway has been shown to turn off, after activation, in response to a constant and ongoing genotoxic challenge. This finding has broad sweeping implications, considering the essential role that ATR plays in checkpoint signaling activation. Loss of effective checkpoint signaling is a hallmark of cancer, promoting genome instability and mutagenesis. The results here show that ATR signaling maintenance is regulated by PARP1 and PAR (Fig. 3). PARP1 activity and protein stability are thus implicated in the ability of cells to activate cell cycle arrest through ATR-Chk1 checkpoint signaling faithfully and may play additional uncharacterized roles as a tumor suppressor gene by connecting metabolic availability of NAD^+^ and DDR signaling. This idea of PARP1 as a tumor suppressor is especially relevant considering the increasing clinical use of PARP1 inhibitors as frontline strategies for treatment of breast and ovarian cancer. It may offer some explanation why these inhibitors have fallen disappointingly short of clinical outcome expectations.

To elucidate how PARP1 promotes DSB formation and ATR-Chk1 signaling after MMS challenge, we examined nuclear extract derived from MMS-treated cell populations. We introduced a defined AP-site substrate into our nuclear extracts and found that MMS treatment reduces exonuclease activity in a temporal-dependent manner (Fig. 4). We also showed that PARP1 and NAD^+^ availability are critical regulators of this exonuclease activity (Fig. 4). Extrapolating to cells, exonuclease activity is capable of shortening the distance constraints between adjacent SSB and increasing intracellular ssDNA content. While technically challenging to prove *in vivo*, the regulation of exonuclease activity in isolated nuclear extract provides a mechanistic explanation for how PARP1 and PAR may regulate DSB formation and ATR-Chk1 activation. We identify two candidate exonucleases which may promote these phenotypes: APE1 and APE2. Inhibitors for these proteins reduced exonuclease activity in our nuclear extracts and reduced DSB formation after MMS challenge (Fig. 4). How and if PARP1 regulates APE1 and APE2 directly or indirectly remains open questions for future study. Due to the complexity of PARP1 and PAR as regulatory factors, we were unable to prove a direct relationship between PARP1, PAR, APE1, and APE2. We performed preliminary experiments using mass spectrometry to identify co-immunoprecipitants with anti-PAR antibodies after MMS challenge. However, these experiments failed to identify APE1 or APE2 (data not shown). We did, interestingly, obtain evidence for KU70 and KU80 PARylation, which had been previously reported^83,84^. While we don’t provide evidence showing that PARP1 directly regulates APE1 and APE2, indirect mechanisms of regulation are equally likely to influence our phenotypes. These mechanisms may include peripheral but important processes such as chromatin remodeling to support exonuclease processing.

Of note, previous work on IR-induced DSB supports a similar SSB-to-DSB conversion model that we propose here, where local oxidative damage induced by IR will result in proximal base damage and require BER for a DSB conversion event^47^. Because this model appears to be conserved in our system, we believe that intracellular levels of NAD^+^ may play a previously uncharacterized role in cell fate, DSB generation, and DDR signaling observed after IR, possibly similar to what we have described here for MMS.

Metabolic disruption is one of the key features of cancer development and progression^85^. A better understanding of metabolic pathways, and key metabolites such as NAD^+^, is critical for understanding the potential efficacy of front-line cancer treatments including IR and chemotherapy. Chemotherapy that results in non-helix distorting base damage, such as alkylating agent N-methyl-N’-nitro-N-nitrosoguanidine (MNNG), may also rely on BER for the generation of downstream toxic repair intermediates^86^. Cancer-specific analysis of these BER factor expression levels may be a key pre-treatment step to evaluate the likelihood of a favorable clinical outcome. For example, MNNG lesions are processed by methylguanine methyltransferase (MGMT), and clinicians usually assess the expression level of MGMT prior to use of MNNG because of the predictive potential of MGMT expression on clinical outcome^77^. Similar strategies may be needed to assess BER protein expression and NAD^+^ levels as a predictive feature of certain chemotherapies.

Taken together, these findings reveal new ideas relating to environmental exposure to DNA-damaging agents, especially chemical modifiers of DNA bases like MMS and Ionizing radiation. The idea that repair intermediates are a source of genome instability themselves has been appreciated for some time^7,87–89^. We support these claims by demonstrating that downstream repair intermediates are responsible for generating lesions that drive DDR activation and DSB formation, rather than the direct chemical activity of MMS. Our findings reproduce some previous work done on MMS, such as MPG’s requirement for DDR activation and DSB formation^9^, but also contributes new ideas such as novel regulatory dynamics of ATR-Chk1 signaling and DSB generation dynamics after MMS exposure which enhances our understanding of how DNA damage is sensed and processed within cells.

### Limitations of the study

Earlier studies have mainly relied on the generation of γH2AX as an indicator of DSB; however, we now know that γH2AX can be produced by multiple types of lesions other than DSB (and hypothetically any cellular condition that activates ATM, ATR, and/or DNA-PKcs can result in γH2AX). This may explain some of the disparity in conclusion between this study and previous studies^9^. We recognize that a similar pitfall may apply to 53BP1, which may complicate our interpretation of the data. Consistent with ours, 53BP1-γH2AX colocalization has been used as a marker or indicator of DSB generation in several studies^90,91^. To improve the rigor of our data, we performed neutral comet assays^54^ to support our use of colocalization of 53BP1-γH2AX foci as a reliable marker for DSB formation and dissolution dynamics. While we think this approach is sufficient to generate convincing evidence for DSB generation analysis, other approaches such as genome-wide DSB mapping via specialized sequencing workflows may provide additional confidence in the DSB generation and resolution model.

Additionally, 3mM MMS is an extremely high dose for cellular exposure. We recognize that this dose may be non-physiological and unlikely to be achieved by traditional chemotherapy agents. Instead, we justify our use of this dose by extrapolating our results to IR, which generates extremely high amounts of local DNA damage (exact dose, time, and tissue depth affect DNA damage loads from IR) in a similar manner to MMS. IR and MMS rely on many of the same genes for sensitivity, so much so that MMS has been historically characterized as a ‘radiomimetic’. The MMS dose used here revealed unique phenotypes that may also occur in lower-dose/longer-exposure situations but be more difficult to detect. We would also like to refer to several previous publications which have used similar concentrations of MMS (0.5-2.5mM)^9,12^. We hope to address these ideas in our future work.

## STAR METHODS

### EXPERIMENTAL MODEL DETAILS

#### Cell culture

Wild type (WT) U2OS (HTB-96) and PANC1 (CRL-1469) cells were purchased from ATCC. MPG-KO and APE1-KO U2OS cells were a generous gift from Dr. David Pellman (HHMI, Dana-Farber cancer institute)^92^. PARP1-KO U2OS cells were a generous gift from Dr. Shan Zha (Columbia University)^93^. All U2OS and PANC1 cells were cultured in Dulbecco’s modified Eagle’s medium (DMEM) supplemented with 10% FBS and penicillin (100 U/ml) and streptomycin (100 μg/ml) at 37°C in 5% CO2 incubator.

### METHOD DETAILS

#### Immunofluorescence Microscopy Analysis

Cells were seeded onto cover slides one day prior to immunofluorescence microscopy analysis. After relevant exposures, cells were washed with PBS and fixed in 4% paraformaldehyde dissolved in phosphate-buffered saline (PBS) for 15 min. After three PBS washes, cells were permeabilized in 0.3% Triton-X dissolved in PBS for 5 min at room temperature. Cells were again washed three times with PBS and then incubated in blocking buffer (5% BSA in PBS) for one hour at 37°C. Cells were incubated with different antibodies diluted 1:100 in blocking buffer at 4°C overnight. If only fluorophore-conjugated primary antibodies were used, samples were washed three times with PBS and directly mounted with ProLong Gold Antifade with DAPI. If unconjugated primary antibodies were used, cells were washed three times with PBS then incubated with appropriate conjugated goat anti-rabbit/mouse secondary antibodies for one hour at 37°C then washed three times with PBS before mounting with ProLong Gold Antifade with DAPI. After mounting, slides cured for one day at room temperature in a dark room. Fluorescent images were acquired using the DMi8 Leica THUNDER Imager without any deconvolution application. All images within each experiment were taken at the same laser and exposure intensity using 63x objective oil-immersion lens. Images formatted for publication which were not used for florescence intensity calculations were exposure adjusted for better readership interpretation (raising exposures can reveal difficult-to-see imaging features such as foci especially if there is a large difference between channel intensities within an experiment (e.g., γH2AX foci after three-hours MMS are much brighter than after one-hour, without raising exposure on one-hour MMS images, clear interpretation of this data would be difficult for readers). No exposure-adjusted images were used for intensity quantifications. Any post-acquisition image adjustment was made to the entire image and not done in any way that may obscure any particular feature of the image. All raw data images are available upon request. Antibodies for immunofluorescent analysis were purchased from respective vendors detailed in the Key Resources Table.

#### COMET Assays

COMET assays were performed using Cell Biolabs Inc OxiSelect COMET Assay Kit (STA-351) following the manufacturer’s protocols. Briefly, cells were treated, trypsonized, and washed with PBS. Cells were resuspended at 1×10^5^ cells/mL before combining the cell suspension with melted (37°C) Comet agarose and subsequent transfer to the relevant well on the Comet slide. After solidification, agarose-cell pucks are incubated in Lysis Buffer (kit) and then incubated with Alkaline Solution (kit). Neutral/TBE conditions involve aspiration of an Alkaline Solution, followed by 15 minutes electrophoresis at 1 volt/cm using cold TBE Electrophoresis Solution (kit). For Alkaline conditions, slides are transferred to an electrophoresis chamber containing Alkaline Electrophoresis Solution (kit), and 1 volt/cm is applied for 15 min. After either Neutral or Alkaline conditions, slides are then washed twice with distilled water and once with 70% Ethanol. After air drying, DNA staining with Vista Green DNA Dye is performed before fluorescent analysis and image acquisition using appropriate filters on the DMi8 Leica THUNDER Imager without any deconvolution application. All images within each experiment were taken at the same laser and exposure intensity. Images were analyzed using Comet Assay IV Lite software.

#### Preparation of Total Cell Lysates and Nuclear Extracts

For stress condition experiments, cells were treated with MMS (Sigma Cat#129925) at 3mM final concentration dissolved in Complete DMEM media, and incubated for the indicated times before cell collection and further analysis. Inhibitors used in this study are as follows: Nicotinamide Riboside Chloride (Selleckchem Cat#S2935), FK866 (Selleckchem Cat#S2799), PDD00017273 (PARGi) (Selleckchem Cat#S8862), Olaparib (Selleckchem Cat#S1060), NU7441 (Selleckchem Cat#S2638), to final concentrations and incubated for times as indicated. Inhibitors were dissolved in DMSO or distilled water as stock solutions and saved at –20°C or -80°C for use per the manufacturer’s recommendation.

After various treatments, cells were washed with PBS and resuspended in Lysis Buffer (20 mM Tris–HCl pH 8.0, 150 mM NaCl, 2 mM EDTA, 0.5% NP-40, 0.5 mM Na3VO4, 5 mM NaF, 5 μg/ml of Aprotinin and 10 μg/ml of Leupeptin). As recently described^68,94^, the total cell lysates were isolated by centrifugation (12,000 rcf for 30 min at 4°C). Nuclear extracts were prepared as previously described^70,95^ and briefly as follows. After PBS washing, cells were resuspended in a suspension buffer (20 mM Tris–HCl pH 7.4, 10 mM NaCl, 3 mM MgCl2) and incubated on ice for 15 min. Then, NP-40 was supplemented to a final concentration of 0.5% and vortexed. The samples were centrifuged (3,000 rpm at 4°C for 10 min) to separate permeabilized nuclei from the cytoplasmic fraction. The recovered nuclei were lysed with Lysis Buffer followed by centrifugation (13,000 rpm at 4°C for 30 min) for nuclear extracts.

#### In-Vitro Nuclease Analysis of Defined AP-site Structure With Nuclear Extracts

For analysis of BER and nuclease activity of nuclear extracts, cells were first treated as indicated prior to nuclear extract isolation. Bradford assays were then performed, and 20μg of nuclear extract was then incubated with 1μg of our annealed double stranded AP-site mimic oligo for the indicated time at 37°C. For Celastrol and AR03 experiments, nuclear extracts were treated with indicated concentrations of inhibitors for 30 min on ice prior to addition of the DNA construct. The AP-site was mimicked via the insertion of a tetrahydrofuran (THF) group at the indicated position, as recently described^95^. Nuclease assay reactions were quenched with equal volume of TBE– urea sample buffer and denatured at 95°C for 5 min. After a quick spin, samples from nuclease assays were examined on 6% TBE–urea PAGE gel and imaged with a Bio-Rad ChemiDoc MP Imaging System. Recombinant His-tagged human APE1 protein (His-APE1) was expressed and purified from E.coli using plasmid pET28HIS-hAPE1, a gift from Primo Schaer (Addgene plasmid #70757; http://n2t.net/addgene:70757; RRID: Addgene 70757)^96^. APE1 protein was used as a positive control.

#### Cell Synchronization & Flow Cytometry Cell Cycle Analysis

For synchronization of cells to G1 phase, we used the established Thymidine-Nocodazole double block synchronization protocol outlined previously^82^. Briefly, 10cm dishes of U2OS cells are arrested in S-phase using a 2mM thymidine incubation for 20h. Cells were washed and released from S-phase thymidine block for 5 hours prior to treatment with 50ng/mL of the microtubule inhibitor Nocodazole. Cells were incubated with Nocodazole for 11 hours and detached via “mitotic shake off”. Mitotic cells are then plated into 6-well dishes for downstream applications and allowed to attach and enter G1 phase for 3 hours prior to drug treatments.

For Flow Cytometry cell cycle analysis, cells were stained with DAPI and resuspended in PBS. 10,000 events were then measured using the BD LSRFortessa flow cytometer and gating was performed using the BD FACSDiva Software considering the roughness and size of events. DAPI staining then indicates the DNA content to allow for quantification of cell cycle distribution. Flow Cytometry cell cycle analysis of synchronized cells was performed in parallel for all experimental conditions to confirm successful G1 phase synchronization. Synchronization escape was typically less than 5% in our experiments and we do not predict this to be a major contributor to our reported phenotypes.

#### Immunoblotting Analysis and Antibodies

Immunoblotting analysis was performed as described previously^70,94^. Briefly, 10μg of cell lysate was loaded onto a 6% Bis-Tris polyacrylamide gel and run at 25mA per gel for 40 minutes. Gels were then transferred onto methanol-activated PVDF membrane for 100 minutes at 100 V. PVDF membranes are then blocked using 5% milk-TBST. Primary antibodies are then diluted 1:000 in 5% BSA-TBST and added to the blocked membrane. Primary antibodies are incubated overnight at 4°C. Primary antibodies are removed, and blots are washed using TBST. Secondary antibodies are diluted 1:2000 in 5% milk-TBST and incubated for 2 hours at room temperature. Membranes are washed again with TBST prior to incubation with WesternBright ECL (Advansta, K-12045). Membranes were then imaged using a Biorad Chemidoc imager. Primary and secondary antibodies were purchased from respective vendors detailed in the Key Resources Table.

#### Cell Viability Analysis

For cell viability analysis using MTT (Thiazolyl blue tetrazolium bromide) assays, cultured cells in a 96-well plate were examined using a procedure as recently described with some slight modifications^70^. Here, we modified the protocol to expose cells to different times of MMS and performed a six-hour recovery prior to reading cell absorbance at 570 nm. To perform the MTT assay, cells were seeded into a 96-well plate at a density of approximately 5,000 to 10,000 cells per well in 100 µL of culture medium. Cells adhered to the plate by incubating it overnight at 37°C in a 5% CO₂ atmosphere. Cells were then treated with the indicated conditions and allowed to recover for six hours. During the last two hours of the recovery period, 10 µL of MTT solution (prepared at 5 mg/mL in PBS) was added to each well. Plates were then incubated for an additional 2 hours at 37°C to allow viable cells to reduce the MTT reagent to insoluble purple formazan crystals. Following incubation, the medium is removed from each well. 150 µL of solubilizing agent was added to each well to dissolve the formazan, and the plates were rocked for about 10 minutes to ensure complete solubilization. Finally, absorbance was measured at 570 nm using a Thermo Scientific Multiskan GO microplate reader.

#### EdU Labelling and Analysis

We labelled cells using ThermoFisher Scientific Click-iT EdU Imaging Kit and protocol. Briefly, cells are incubated on coverslips overnight before being incubated with 10μM EdU for 3 hours. MMS treatment occurs in the final hour of incubation as indicated. Cells are fixed in 4% paraformaldehyde for 15 minutes at room temperature, then washed with PBS, and permeabilized with 0.3% Triton X-100 in PBS at room temperature for additional 15 minutes. Primary antibody is then added and incubated overnight. After washing, Click-iT reaction cocktail is then added to coverslips (manufacturer recipe) and incubated for 30 minutes in the dark. Cover slips are then mounted and imaged as described above. For quantification of cells, EdU labelled cells fluoresce green as S-phase cells, while cells with DAPI positive and EdU negative indicate cells in G1 and G2 phases. 53BP1 foci are counted, and the cell cycle is indicated.

### QUANTIFICATION AND STATISTICAL ANALYSIS

#### Quantification, Statistical analysis, and Reproducibility

The data presented are representative of three biological replicates unless otherwise specified. All immunofluorescent-based foci counting experiments use n=50 cells unless otherwise indicated. All statistics were performed using GraphPad Prism version 8 for Mac. Statistical significance was ascertained between individual samples using a parametric unpaired t-test for immunoblotting, and a nonparametric Kolmogorov-Smirnov test for immunofluorescent analysis data, including COMET and 53BP1-γH2AX foci data. Significance is denoted by asterisks in each figure: *P < 0.05; **P < 0.01; ***P < 0.001; ****P < 0.0001; ns, no significance. Bars represent means and error bars represent the standard deviation (SD) for three independent experiments/n=50 cells, unless otherwise indicated.

### KEY RESOURCES TABLE

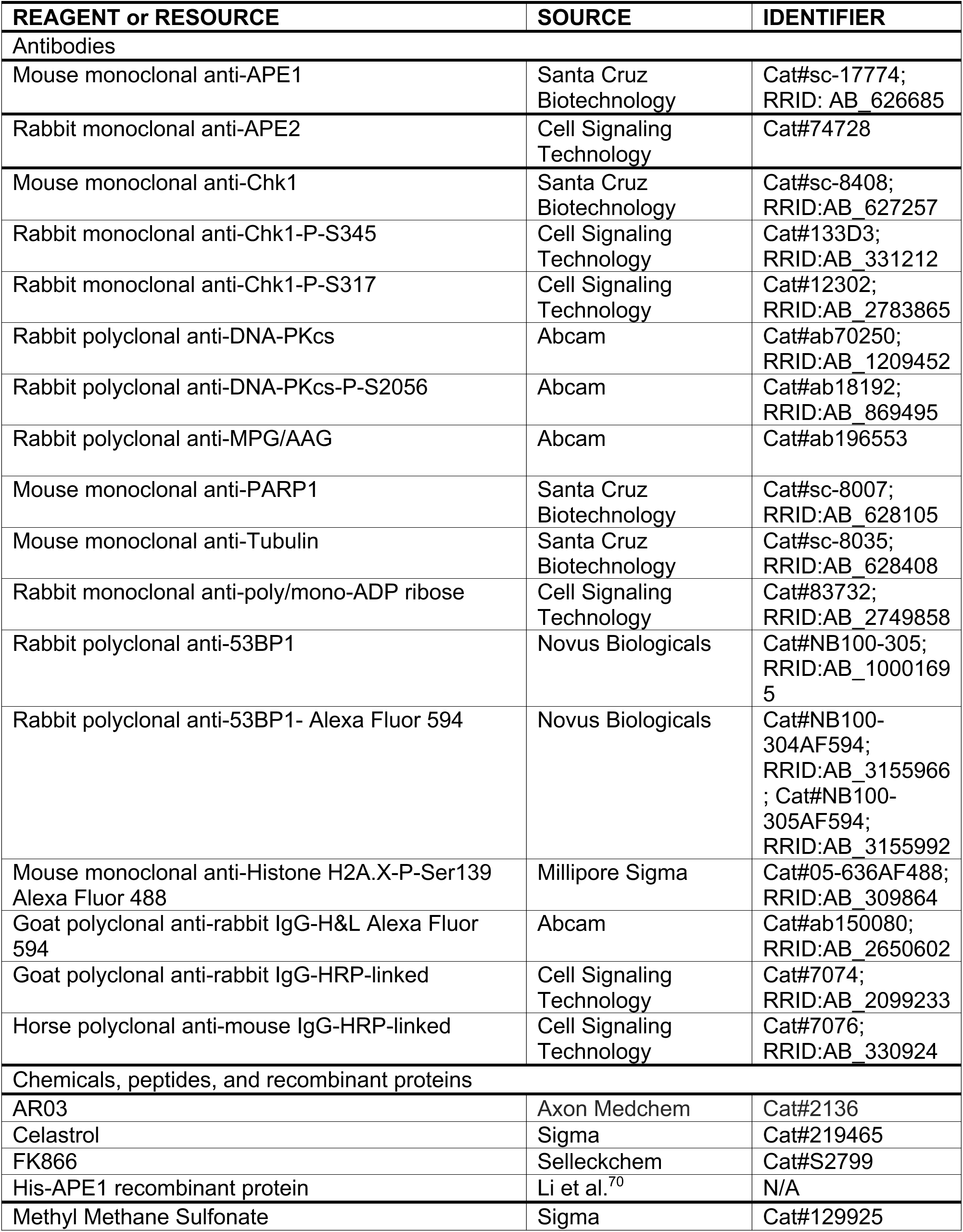
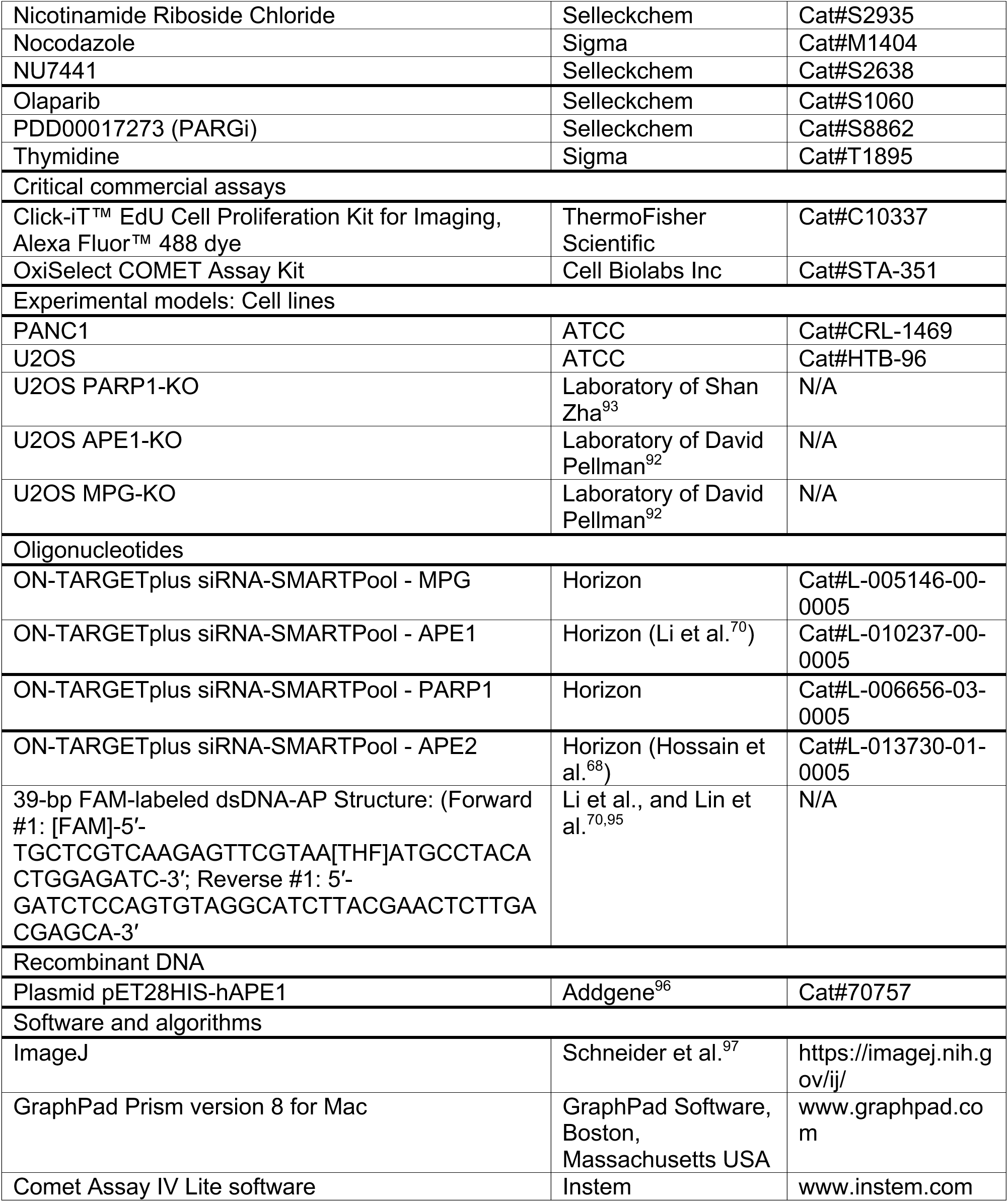

## ACKNOWLEDGEMENTS

We are grateful to Dr. Primo Schaer, Dr. David Pellman, and Dr. Shan Zha for reagents and materials. The Yan lab is supported in part by the funds from University of North Carolina at Charlotte and grants from the NIH/NIEHS (R21ES032966 to S.Y.) and the NIH/NCI (R01CA291546, R01CA225637 and R03CA270663 to S.Y.). A.M.M., G.D., J.M., were supported in part by UNC Charlotte’s Graduate School Summer Fellowship program. J. Lyttle was supported in part by the OUR Scholars program at UNC Charlotte. S.Y. thanks Zhe Li for reading and comments on the manuscript.

## AUTHOR CONTRIBUTIONS

Conceptualization: A.M.M., S.Y.; Formal analysis: A.M.M., H.Z., S.Y.; Funding acquisition: S.Y.; Investigation: A.M.M., H.Z., J.Li, G.D., J.M., K.M., J.Lyttle, S.Y.; Methodology: A.M.M., H.Z., J.Li, H.Z., J.Lyttle, G.D., J.M., K.M., J.L., S.Y.; Project administration: S.Y.; Supervision: S.Y.; Visualization: A.M.M., H.Z., S.Y.; Writing – original draft: A.M.M., S.Y.; Writing – review & editing: A.M.M., S.Y.

## ADDITIONAL INFORMATION

### Supplementary information

The online version contains five supplementary figures (Figs. S1 to S6).

**Correspondence** and requests for materials should be addressed to Shan Yan

## Figures and Figure Legend

**Figure S1.**
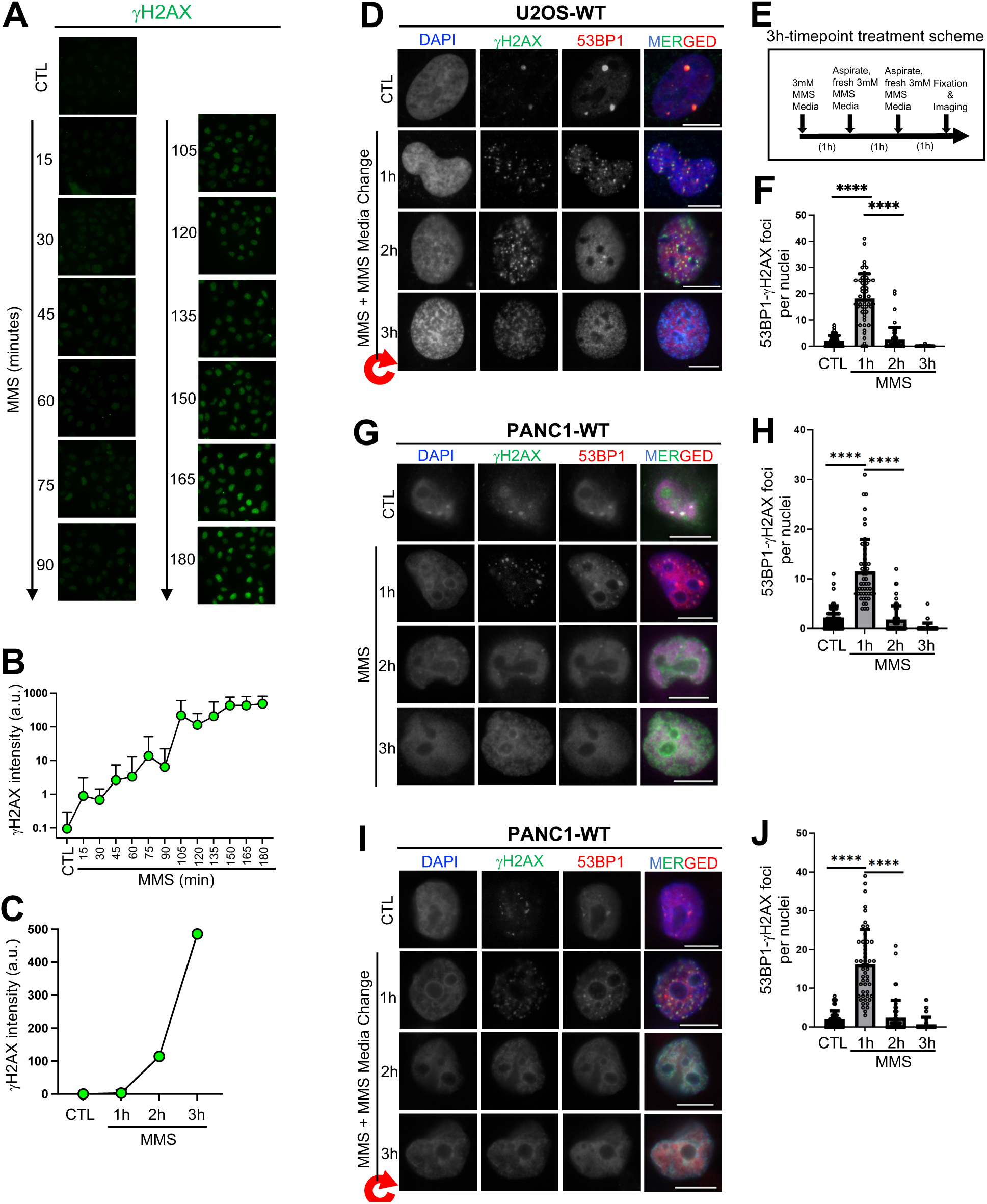
γH2AX intensity analysis and colocalizing γH2AX-53BP1 foci analysis in PANC1 cells following different MMS treatment (Related to Figure 1). *A*, γH2AX intensity was measured after 15-minute time course exposure to 3mM MMS in WT U2OS cells. *B-C*, γH2AX intensity was quantified from *A* using ImageJ. *D*, Immunofluorescent microscopy analysis was performed using DAPI (nuclei staining), γH2AX (AF488), and 53BP1 (AF594) after exposing U2OS cells to 3mM MMS-containing media and changing the media every hour. *E*, Sample treatment scheme for three-hour MMS treated samples. *F*, Overlapping/colocalizing γH2AX-53BP1 foci were quantified on a per nuclei basis using n=50 cells after media swap condition in *D*. *G*, PANC1 cells were used for immunofluorescent microscopy using DAPI (nuclei staining), γH2AX (AF488), and 53BP1 (AF594) after exposing cells to one, two, and three hours of 3mM MMS respectively. *H*, Overlapping/colocalizing γH2AX-53BP1 foci were quantified on a per nuclei basis using n=50 cells. *I*, PANC1 cells were used for immunofluorescent microscopy analysis using DAPI (nuclei staining), γH2AX (AF488), and 53BP1 (AF594) after exposing the cells to 3mM MMS-containing media and changing the media every hour. *J*, Overlapping/colocalizing γH2AX-53BP1 foci were quantified on a per nuclei basis using n=50 cells for the media swap experiment in *I*. For all experiments, significance was determined using the Kolmogorov-Smirnov test and is demonstrated as follows: ****, p<0.0001. Scale bar = 10μm.

**Figure S2.**
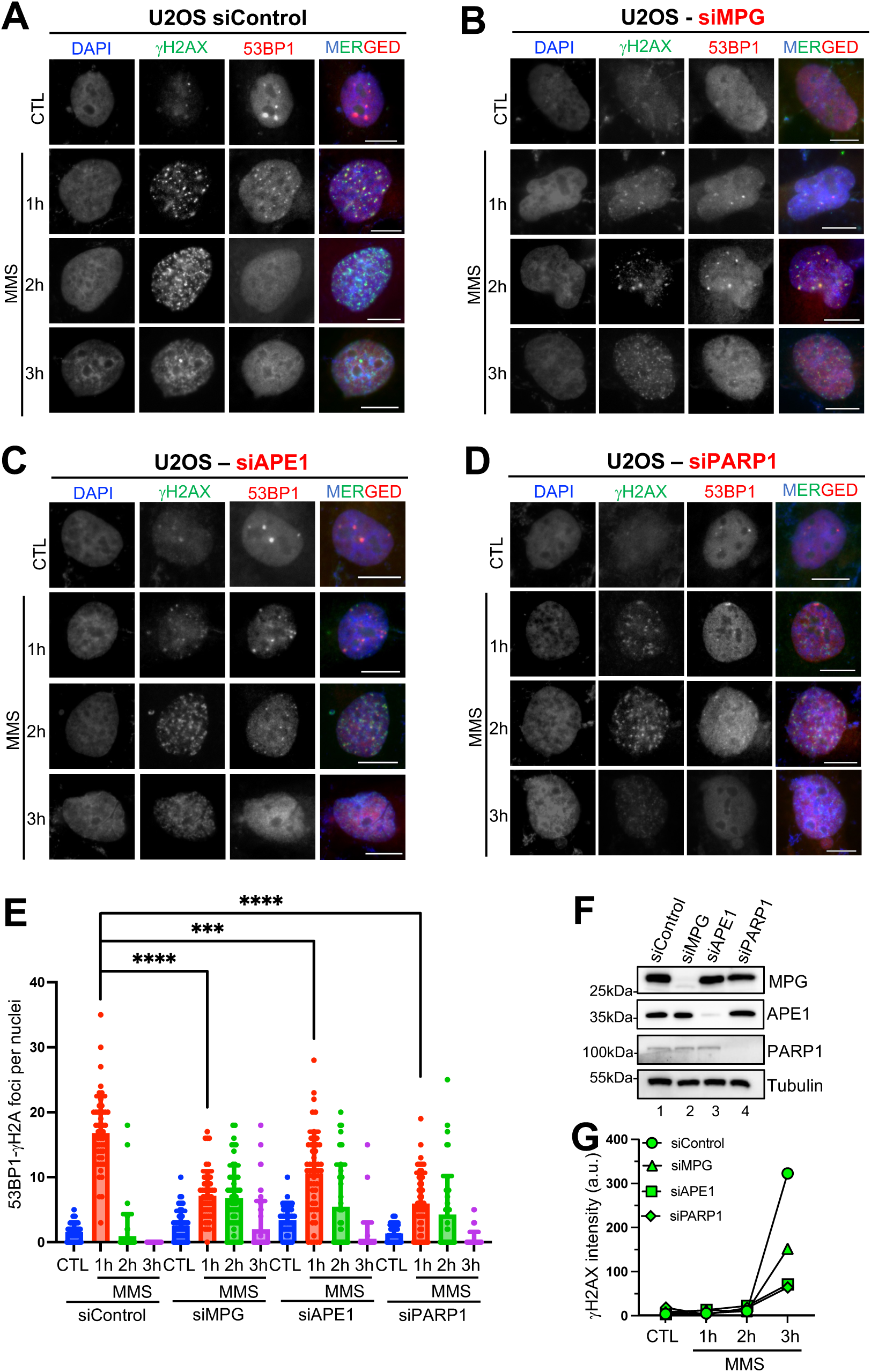
γH2AX intensity analysis and colocalizing γH2AX-53BP1 foci analysis in U2OS cells in control or target gene was knocked down (Related to Figure 1). *A*, siRNA control was performed on U2OS WT cells prior to immunofluorescent microscopy using DAPI (nuclei staining), γH2AX (AF488), and 53BP1 (AF594) after exposing cells to one, two, and three hours of 3mM MMS, respectively. *B*, siRNA against MPG was performed on U2OS WT cells prior to immunofluorescent microscopy using DAPI (nuclei staining), γH2AX (AF488), and 53BP1 (AF594) after exposing cells to one, two, and three hours of 3mM MMS, respectively. *C*, siRNA against APE1 was performed on U2OS WT cells prior to immunofluorescent microscopy using DAPI (nuclei staining), γH2AX (AF488), and 53BP1 (AF594) after exposing cells to one, two, and three hours of 3mM MMS respectively. *D*, siRNA against PARP1 was performed on U2OS WT cells prior to immunofluorescent microscopy using DAPI (nuclei staining), γH2AX (AF488), and 53BP1 (AF594) after exposing cells to one, two, and three hours of 3mM MMS, respectively. *E*, Quantification of parallel siRNA experiments using n=50 cells per condition and Kolmogorov-Smirnov test to determine statistical significance. *F*, Western blot confirmation of siRNA knockdown performed in parallel with imaging experiments. *G*, ImageJ quantification of γH2AX intensity between different siRNA-treated groups. For all experiments, statistical significance is demonstrated as follows: ***, p<0.001; ****, p<0.0001. Scale bar = 10μm.

**Figure S3.**
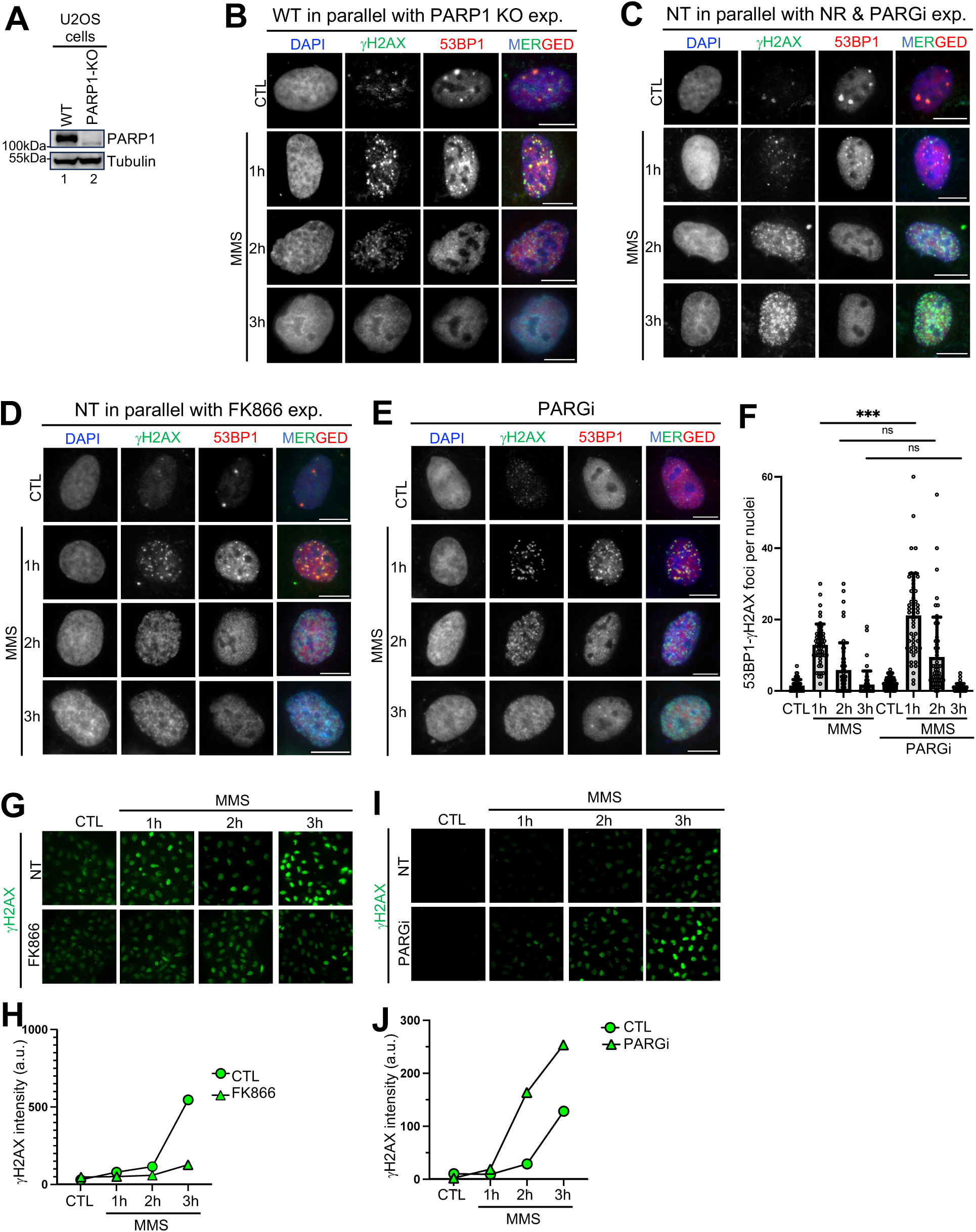
γH2AX intensity analysis and colocalizing γH2AX-53BP1 foci analysis in U2OS cells under different treatments (Related to Figure 2). *A*, Immunoblot confirmation of PARP1-KO cells was performed in parallel with WT U2OS cells. *B-D*, U2OS WT were imaged using DAPI (nuclei staining), γH2AX (AF488), and 53BP1 (AF594) after exposing cells to one, two, and three hours of 3mM MMS, respectively, in parallel with PARP1-KO, NR & PARGi, and FK866 conditions, respectively. *E-F*, PARGi treatment was performed before immunofluorescent microscopy using DAPI (nuclei staining), γH2AX (AF488), and 53BP1 (AF594) after exposing cells to one, two, and three hours of 3mM MMS, respectively. Overlapping/colocalizing γH2AX-53BP1 foci were quantified on a per nuclei basis using n=50 cells. *G-H*, γH2AX intensity was quantified using ImageJ, using parallel experiments with and without FK866 treatment (50μM). *I-J*, γH2AX intensity was quantified using ImageJ, using parallel experiments with and without PARGi treatment (20μM for two hours pre-treatment). For all experiments, significance was determined using the Kolmogorov-Smirnov test and is demonstrated as follows: ***, p<0.001; ns, no significance. Scale bar = 10μm.

**Figure S4.**
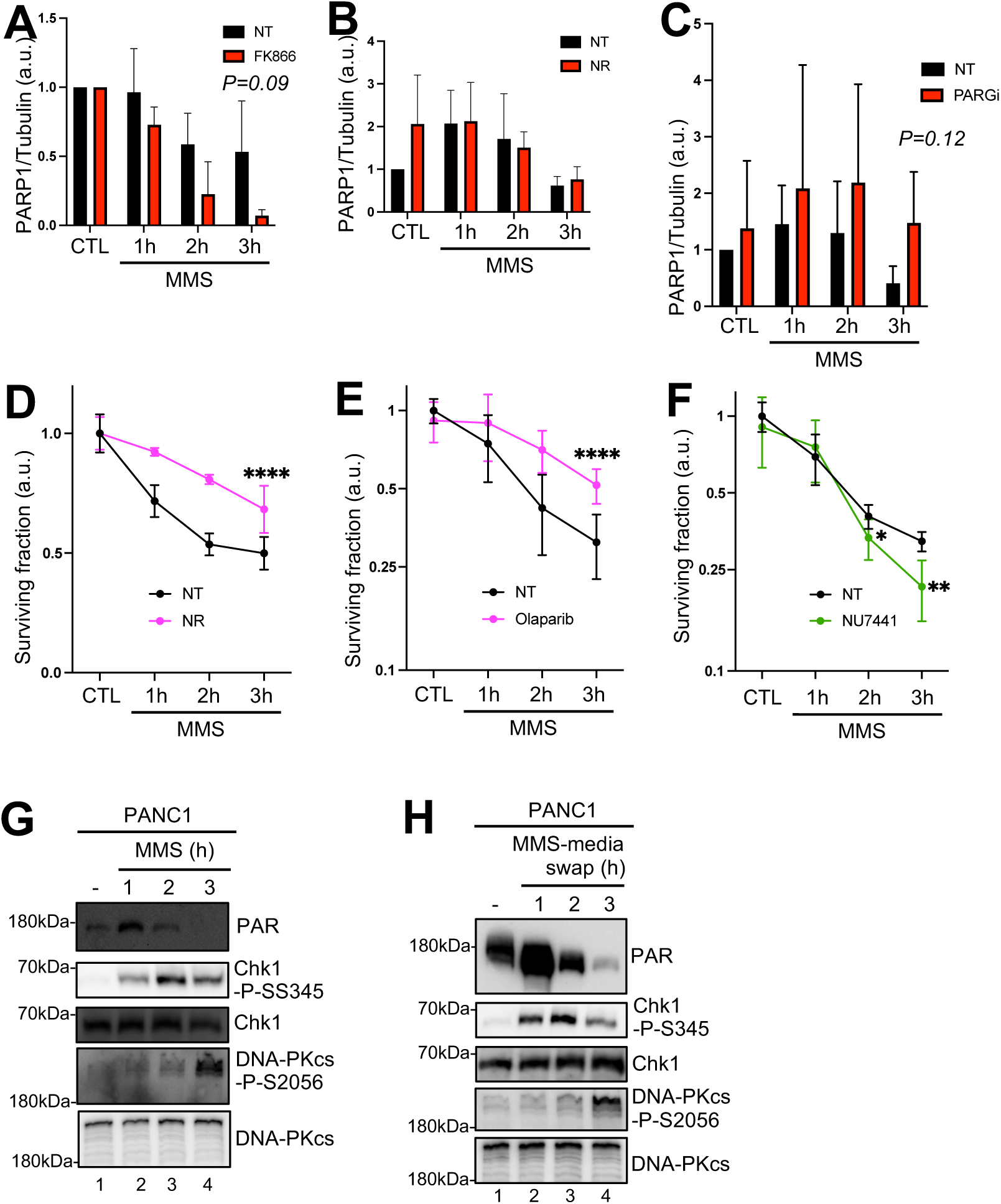
PARP1 abundance and cell survival analysis as well as PANC1 cell analysis (Related to Figure 3). *A*, PARP1 protein abundance was quantified from three biologically independent experiments and normalized to Tubulin after FK866 pre-treatment (50μM for 24 hours pre-treatment) and exposure to 3mM MMS for the indicated timepoints. *B*, PARP1 protein abundance was quantified from three biologically independent experiments and normalized to Tubulin after NR pre-treatment (200μM for 24 hours pre-treatment) and exposure to 3mM MMS for the indicated timepoints. *C*, PARP1 protein abundance was quantified from three biologically independent experiments and normalized to Tubulin after PARGi pre-treatment (20μM for two hours pre-treatment) and exposure to 3mM MMS for the indicated timepoints. *D*, Cell viability was measured via MTT assay after indicated MMS exposure with and without NR supplement (200μM for 24 hours pre-treatment). *E*, Cell viability was measured via MTT assay after indicated MMS exposure with and without Olaparib pre-treatment (10μM for two hours) *F*, Cell viability was measured via MTT assay after indicated MMS exposure with and without NU7441 pre-treatment (10μM for two hours). *G*, PANC1 cells were immunoblotted after MMS exposure, for a total of three-hours of continuous 3mM MMS exposure. *H*, PANC1 cells were immunoblotted after MMS exposure with an MMS-containing media swap every hour, for a total of three-hours continuous MMS exposure. For all experiments, significance was demonstrated as follows: ****, p<0.0001.

**Figure S5.**
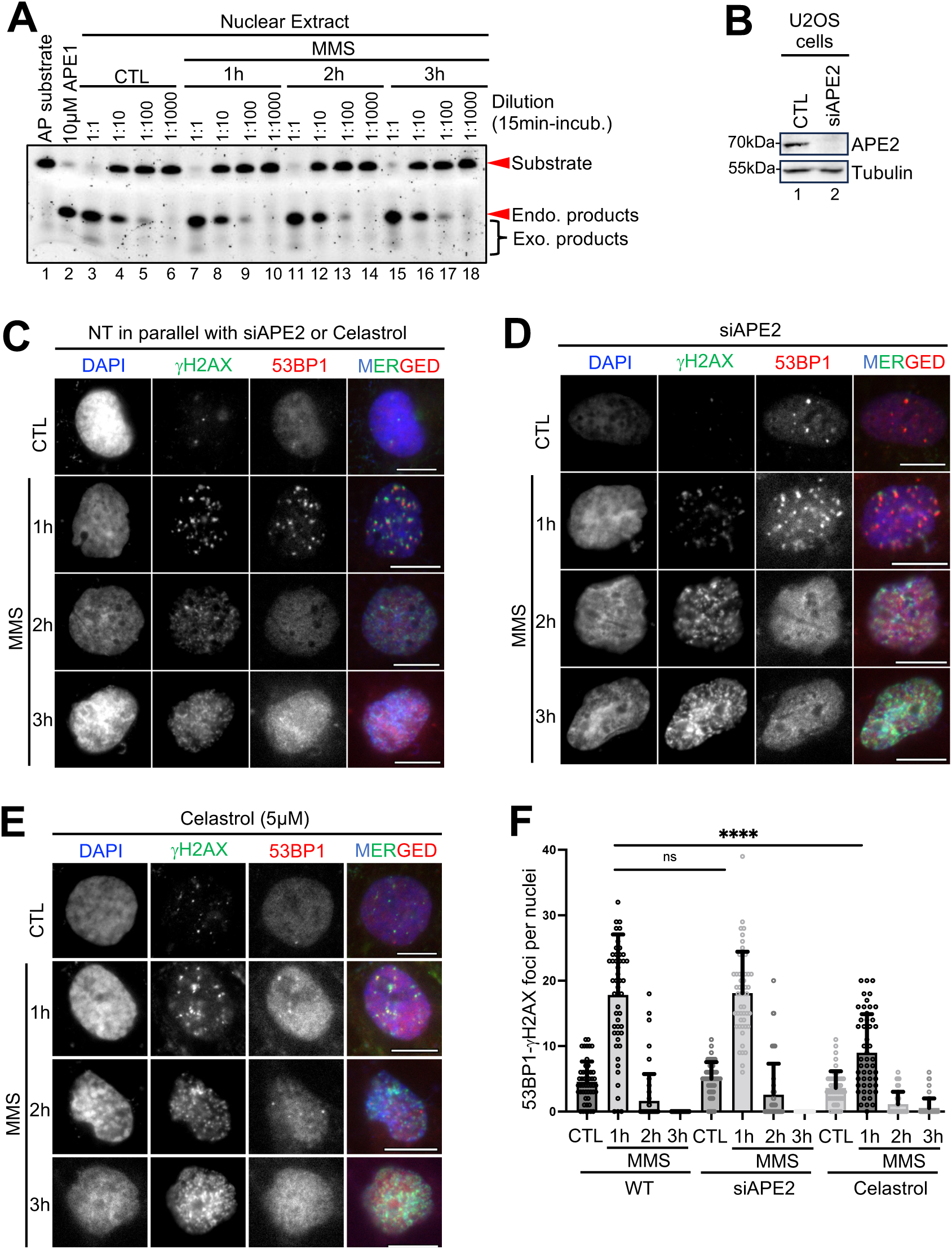
Nuclease activity analysis of AP-mimetic substrate in nuclear extracts and effect of APE2 impairment by siRNA-mediated knockdown or APE2 inhibition by Celastrol in MMS-induced DSB formation (Related to Figure 4). *A*, Nuclear extract was prepared from cells after MMS exposure for indicated times. 20μg of nuclear extract was incubated with the DNA construct (1:1) and then serially diluted before a fifteen-minute incubation at 37°C followed by denaturing EMSA. *B,* Immunoblot confirmation of APE2-KD cells was performed in parallel with CTL-KD cells. *C-F*, siRNA-mediated APE2-KD, Celastrol treatment (5μM for two hours pre-treatment), and siRNA control was performed in parallel prior to imaging using DAPI (nuclei staining), γH2AX (AF488), and 53BP1 (AF594) after exposing cells to one, two, and three hours of 3mM MMS, respectively. Overlapping/colocalizing γH2AX-53BP1 foci were quantified on a per nuclei basis using n=50 cells. The Kolmogorov-Smirnov test was used to determine statistical significance. For all experiments, statistical significance is demonstrated as follows: ****, p<0.0001; ns, no significance. Scale bar = 10μm.

**Figure S6.**
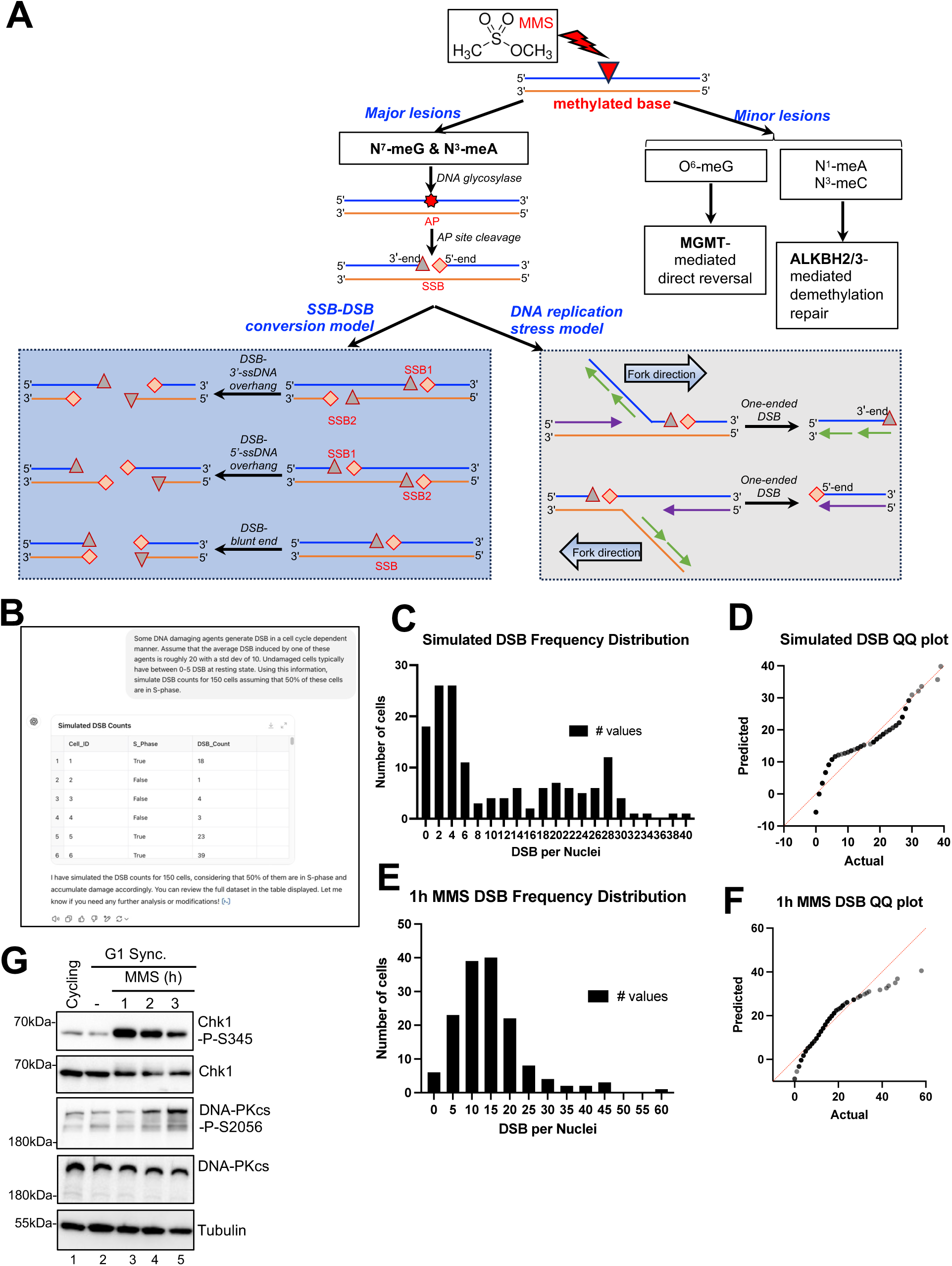
Simulation and validation analysis of MMS-induced DSB frequency distribution in cells (Related to Figure 5). *A*, Conceptual models of MMS-induced DNA damage sensing and processing and DNA repair. B, Screenshot of the prompt used to generate simulated DSB counts. C, A frequency distribution plot was generated using simulated counts for n=150. *D*, QQ Normality plot was generated using simulated counts for n=150. *E*, Frequency distribution plot was generated using γH2AX-53BP1 colocalized foci counts after one-hour 3mM MMS exposure for n=150 cells from three biologically independent replicates. *F*, QQ Normality plot was generated using γH2AX-53BP1 colocalized foci counts after one-hour 3mM MMS exposure for n=150 cells from three biologically independent replicates. *G*, U2OS WT cells were synchronized to G1 phase prior to MMS exposure and immunoblotting.

